# Manifestations of genetic risk for Alzheimer’s Disease in the blood: a cross-sectional multi-omic analysis in healthy adults aged 18-90+

**DOI:** 10.1101/2021.03.26.437267

**Authors:** Laura Heath, John C. Earls, Andrew T. Magis, Sergey A. Kornilov, Jennifer C. Lovejoy, Cory C. Funk, Noa Rappaport, Benjamin A. Logsdon, Lara M. Mangravite, Brian W. Kunkle, Eden R. Martin, Adam C. Naj, Nilüfer Ertekin-Taner, Todd E. Golde, Leroy Hood, Nathan D. Price, Alzheimer’s Disease Genetics Consortium

## Abstract

Deeply phenotyped cohort data can elucidate differences associated with genetic risk for common complex diseases across an age spectrum. Previous work has identified genetic variants associated with Alzheimer’s disease (AD) risk from large-scale genome-wide association study meta-analyses. To explore effects of known AD-risk variants, we performed a phenome-wide association study on ~2000 clinical, proteomic, and metabolic blood-based analytes obtained from 2,831 cognitively normal adult clients of a consumer-based scientific wellness company. Results uncovered statistically significant SNP-analyte associations for five genetic variants after correction for multiple testing (for SNPs in or near *NYAP1, ABCA7, INPP5D*, and *APOE*). These effects were detectable from early adulthood. Sex modified the effects of four genetic variants, with multiple interrelated immune-modulating effects associated with the *PICALM* variant. Sex-stratified GWAS results from an independent AD case-control meta-analysis supported sexspecific disease effects of the *PICALM* variant, highlighting the importance of sex as a biological variable. These analyses support evidence from previous functional genomics studies in the identification of a causal variant within the *PILRA* gene. Taken together, this study highlights clues to the earliest effects of AD genetic risk variants in individuals where disease symptoms have not (yet) arisen.

## Introduction

The rapidly decreasing cost of genomics paired with technological advances in the generation of longitudinal multi-omic data has resulted in multiple datasets of deeply phenotyped individuals with a variety of health outcomes^1–3^. The data collected in these studies have the potential to yield important insights into potential molecular drivers of health, differences observed as a function of differential genetic disease risk, and the earliest manifestations of disease transitions. The present study seeks to leverage a unique and relatively large set of multi-omic, deep-phenotyping data to shed light on genetic pathways to late-onset Alzheimer’s disease (AD) by assessing differences in ~2000 analytes in the blood that show association with known genetic risk variants for AD. Coupled with high-dimensional data sets, this approach has the potential to yield clues into disease processes and possible early-intervention strategies, which are critically important given the essentially untreatable nature of late-stage Alzheimer’s disease once significant brain deterioration has occurred.

Genetic variation plays a substantial role in AD risk, with twin studies estimating AD heritability at 58-79%^4^. While the emergence of recent large-scale consortia efforts has facilitated well-powered meta-analyses of genome-wide association studies (GWAS) to identify multiple common variants with small effect sizes^5,6^, the research community is still untangling exactly how this genetic variation influences disease risk. Functional genomics studies are beginning to identify likely genetic pathways to disease with the aid of transcriptomic, epigenomic, and endophenotypic data^7–10^. So far, genetic and multi-omic studies of AD studies have largely focused on older individuals with either clinically diagnosed AD or milder symptoms of cognitive decline, despite research pointing to highly variable AD pathobiology that occurs on a spectrum, and begins decades before clinical symptoms onset^11^.

In this study, we leveraged the results from a large-scale GWAS meta-analysis^5^ alongside data from a deeply phenotyped wellness cohort to investigate the effects of genetic risk for AD in individuals without cognitive impairment, at all ages. We undertook an agnostic approach by adopting a phenome-wide association study (PheWAS) design^12^. By examining how genetic variation impacts 2008 analytes in the blood of 2831 individuals, we sought to complement previous functional genomics studies as well as potentially reveal new testable hypotheses for future studies. In addition, we tested for associations between a polygenic risk score (PGRS) for AD and blood analytes to determine if a relative burden of genetic risk might impact observable changes in the blood, and we assessed for effect modification of genetic risk by sex.

## Results

### Summary of population and study design

Data was collected from subscribers in a now-closed scientific wellness program (Arivale, Inc.^3,13^), starting in July 2015 and ending in May 2019. From this population, we identified 2831 participants with whole genome sequencing (WGS) and at least one class of blood-derived analyte (clinical chemistries, proteins, or metabolites) obtained at entry into the program (see Online Methods). Sixty-one percent of Arivale participants were female, 22% were of non-white self-reported ethnicity, and 28% were obese (**Table 1**). The mean age at blood draw was 47 years, with a range of 18 to 89+. In general, individuals who joined Arivale had somewhat higher levels of cardiovascular risk markers compared to the US population, and slightly lower rates of obesity and pre-diabetes^3^ (these rates were consistent with rates in the geographies and ethnicities of the population, mostly from the west coast region of the United States).

**Table 1.** Baseline self-reported characteristics of Arivale participants with available whole-genome sequences.

| **Characteristic<sup>a</sup>** | **N=2831** |
| --- | --- |
| Age, mean (sd) | 47.0 (12.0) |
| Female, n (%) | 1719 (60.7) |
| Nonwhite<sup>b</sup>, n (%) (n=2725) | 597 (21.9) |
| Afro-Caribbean | 1 (<0.1) |
| American Indian or Alaska Native | 5 (0.2) |
| Ashkenazi Jewish | 49 (1.8) |
| Asian | 84 (3.1) |
| Black or African American | 64 (2.3) |
| East Asian | 91 (3.3) |
| Hispanic Latino or Spanish origin | 120 (4.4) |
| Middle Eastern or North African | 18 (0.7) |
| Native Hawaiian or other Pacific Islander | 17 (0.6) |
| Sephardic Jewish | 4 (0.1) |
| South Asian | 79 (2.9) |
| White | 2128 (78.1) |
| Other | 65 (2.4) |
| BMI, mean (sd) (n=2750) | 27.9 (6.4) |
| Obese<sup>c</sup>, n (%) (n=2750) | 802 (29.2) |
| Moderate activity $\geq 3$x/wk, n (%) (n=2275) | 1460 (64.2) |
| Vigorous activity $\geq 3$x/wk, n (%) (n=2271) | 697 (30.7) |
| Ever smoke, n (%) (n=2207) | 565 (25.6) |
| Current meds for cholesterol, n (%) (n=2378) | 287 (12.1) |
| Past and/or current self-report of: |  |
| Migraine, n (%) (n=2229) | 558 (25.0) |
| High cholesterol, n (%) (n=2301) | 558 (24.2) |
| Depression, n (%) (n=2278) | 521 (22.9) |
| GERD, n (%) (n=2220) | 464 (20.9) |
| Hypertension, n (%) (n=2316) | 434 (18.7) |
| Asthma, n (%) (n=2361) | 376 (15.9) |

<sup>a</sup>For categories with missing data, total non-missing N is reported in parentheses
<sup>b</sup>Race/ethnicity categories presented to participants in Arivale questionnaire
<sup>c</sup>Obese defined as BMI ≥ 30

We selected 25 common (>5% minor allele frequency (MAF)) and somewhat rare (>1% MAF) single nucleotide polymorphisms (SNPs) linked to 24 genes that were significantly associated with AD in a recent large-scale meta-analysis^5^ (**Supplementary Table 1**). In brief, following a PheWAS approach^14^, for each SNP we fitted a linear regression, with genotype as the independent variable and each quantitative log-transformed analyte as the dependent variable, adjusted for age, sex, vendor (in the case of clinical lab values), and four principal genetic components to account for ancestry. We applied the Benjamini-Hochberg false-discovery rate (FDR) correction^15^ to account for multiple comparisons. We also assessed for gene by sex interactions. An *APOE-free* polygenic risk score (PGRS) was also incorporated into the PheWAS model. Lastly, since the SNP candidates for this study were derived from meta-analysis of non-Hispanic white (NHW) cohort populations^5^, we stratified PheWAS by self-identified race (white vs. other) in order to determine whether the statistically significant signals we observed were a result of the majority-NHW Arivale cohort demographic makeup.

### Phenome-wide association study results

We observed 33 SNP-analyte associations that were statistically significant at FDR-adjusted p-value<0.05, with the majority of the associations observed for the *APOE* SNPs (rs7412, or the e2-defining allele, and rs429358, or the e4-defining allele). The other SNPs showing significant associations with at least one clinical chemistry, protein, or metabolite were rs10933431, rs12539172, and rs3752246 (**Figure 1**, **Table S1**).

**Figure 1.**
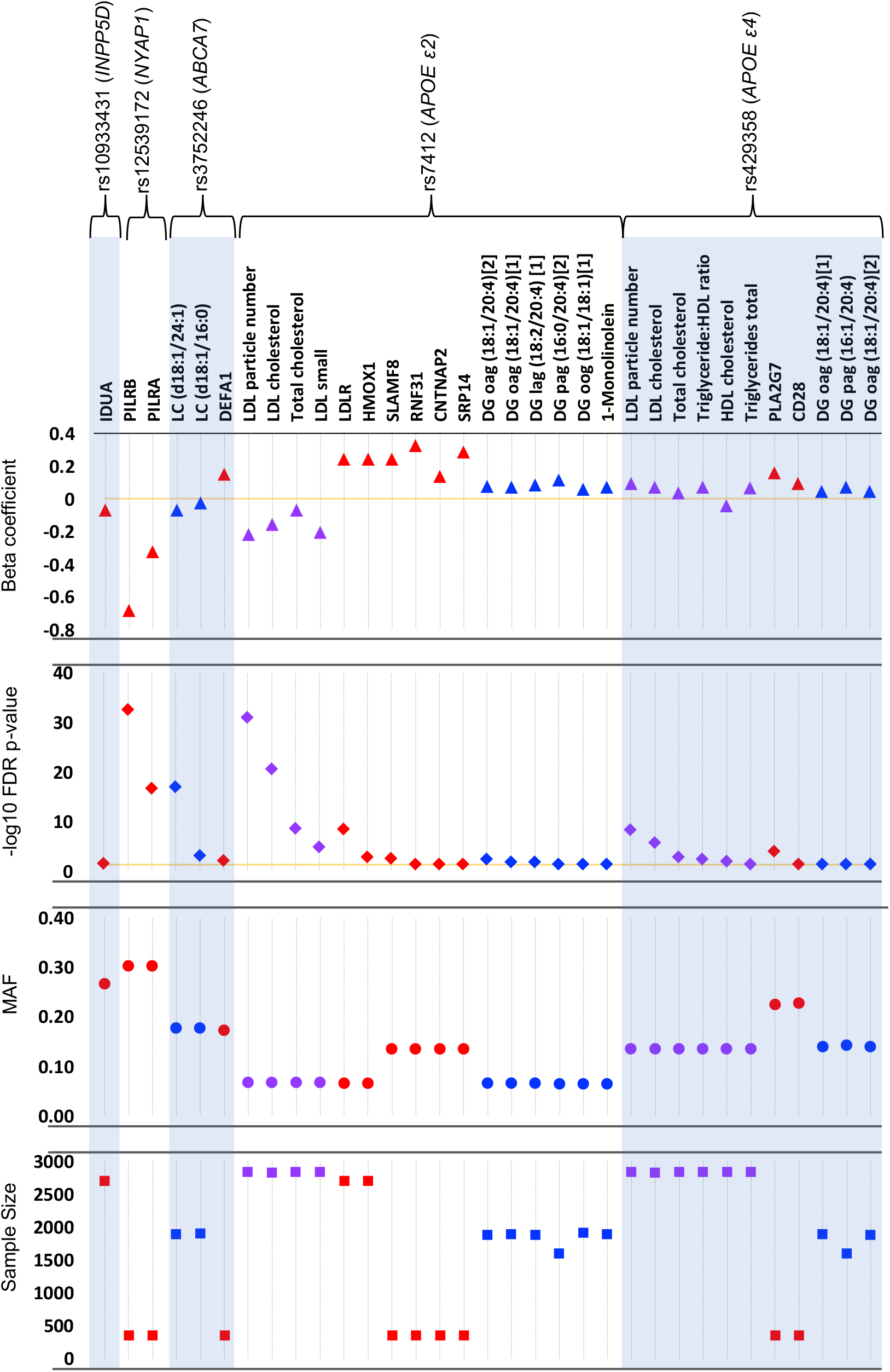
Statistically significant SNP-analyte associations after correcting for multiple testing (threshold FDR-adjusted p-value=0.05), by SNP. Top panel: log-transformed betacoefficient from the linear regression model adjusted for sex, age, and genetic principal components 1-4; markers above the zero line (orange) indicate analytes that increased in value with the minor allele, while markers below the line indicate markers that decreased in value. Second panel: FDR-adjusted −log_10_ p-value; orange line at FDR-p=0.05. Proteins=red, metabolites=blue, clinical chemistries=purple. Metabolite codes: DG=diacylglycerol; LC=lactosylceramide; o=oleoyl; a=arachidonoyl; g=glycerol; l=linoleoyl; p=palmitoyl. Third panel: minor allele frequency (MAF). Bottom panel: Total sample size for each analyte-SNP regression.

#### NYAP1

The most robust SNP-analyte associations we observed were between rs12539712 in the 3’ region of NYAP1, and two co-regulated proteins, paired immunoglobulin-like type 2 receptors beta and alpha (PILRB and PILRA). Carriage of the minor allele (AD risk odds ratio (OR)=0.92) was associated with significant reduction in normalized protein expression (NPX) of PILRB and PILRA compared to individuals homozygous for the major allele (FDR-adjusted p-values=2.2×10^−33^ and 2.3×10^−17^, respectively), while the overall level of NPX increased with age among all genotypes. The reduction in protein levels appears roughly dose-dependent with the number of minor alleles and was observed in all age groups (**Figure 2a**). This locus was originally identified by rs1476679 near *ZCWCP1^6^. NYAP1* and *ZCWPW1* are located near PILRA and PILRB on chromosome 7, within a linkage disequilibrium (LD) block. In previous gene expression studies, the initial index SNP for *ZWCWP1* has been associated with expression of multiple PILRB and PILRA transcripts in brain^9,16^. Other recent studies have pointed to variation in *PILRA* as the causal gene at this locus, with a missense SNP as a leading candidate (G78R, rs1859788)^17–20^. We repeated the PheWAS with this putative causal SNP (which was in LD with our index SNP rs12539172, R^2^=0.77), and the associations became stronger (FDR-adjusted p-value for PILRB=3.6×10^−52^; for PILRA=1.4×10^−22^) (**Figure 2b**).

**Figure 2.**
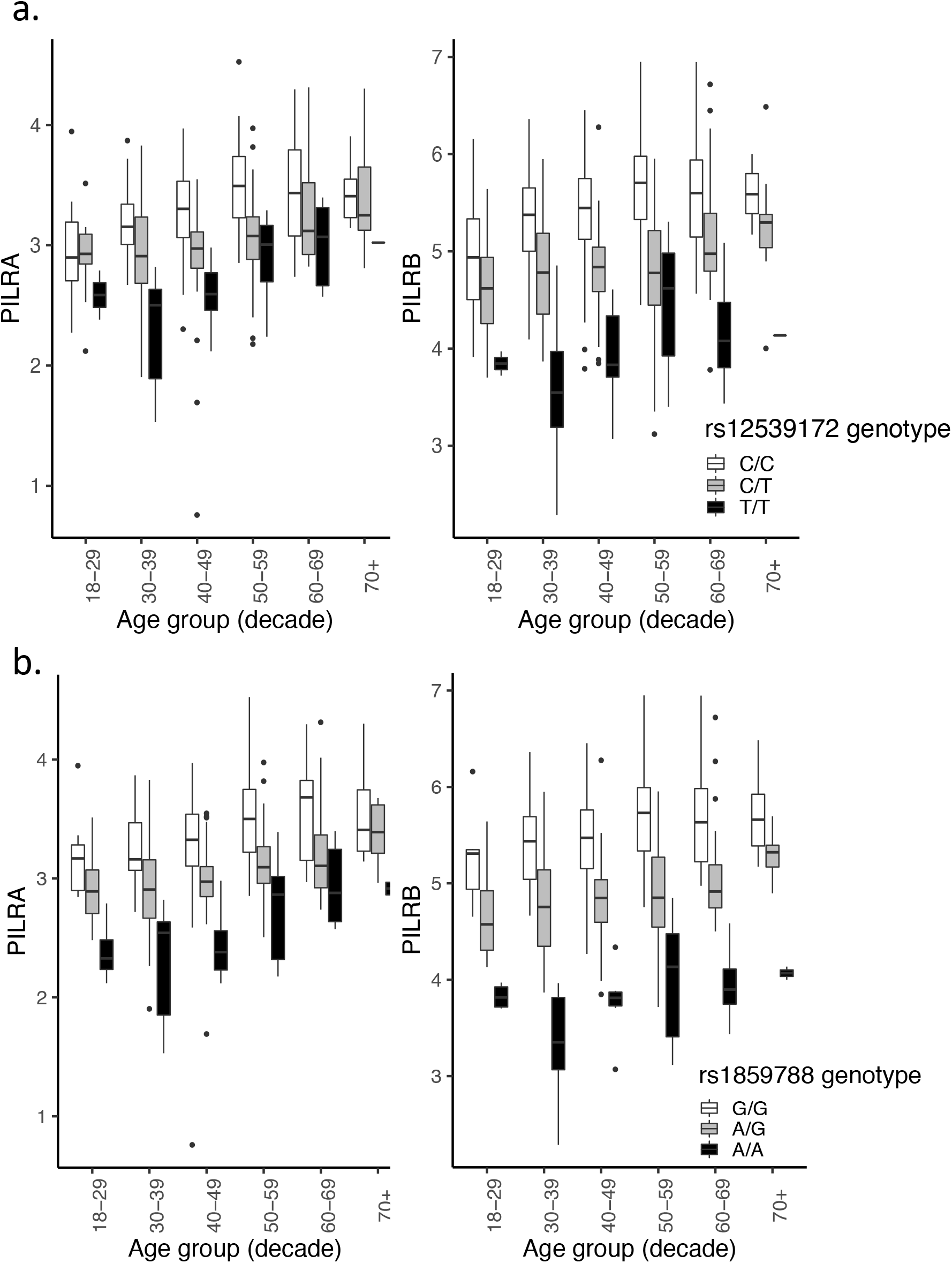
Unadjusted box plots of normalized protein expression (NPX) levels of PILRA and PILRB by genotype and age group. White boxplots=individuals who are homozygous for the major allele, gray boxplots=heterozygotes, black boxplots=minor allele homozygotes. Box plot midline=median value, lower/upper hinges=25^th^ and 75^th^ percentiles, respectively; lower whisker ends/upper whisker ends no further than 1.5 x interquartile range from the hinge. Data beyond whiskers are outlying points. **a.** NPX of PILRA and PILRB by rs12539172 (NYAP1) genotype; **b.** NPX of PILRA and PILRB by rs1859788 genotype.

#### APOE4

We observed significant associations between rs429358 (which encodes the e4 allele) and multiple related clinical measures of cholesterol (**Figure S1**), consistent with previous cohort studies that included young, middle-aged, and older adults^21–24^. Differences by genotype were less pronounced in older age groups likely due to statin use; exploratory analyses visualizing only individuals who did not report use of statin-lowering medications showed more consistent genotype-dependent differences in older age groups (**Figure S2**). The concentration of two inflammation-related proteins in the blood were associated with the e4 allele: PLA2G7 and CD28 (**Figure S1**). PLA2G7 (platelet activating factor acetylhydrolase) is a known cardiovascular risk marker with pro-inflammatory and oxidative activities^25^ which has previously been associated with APOE genotypes^26^ and implicated in AD and cognitive decline^25,27^. Selected lipid metabolites in the blood were positively associated with e4: two diacylglycerol (DG) metabolites (one of which was measured twice in the Metabolon panel) were higher in e4 carriers compared to non-carriers. These DGs were also elevated in *APOE* e2 carriers (see below).

#### APOE2

Consistent with previous studies^21–24^, we observed significantly lower levels of multiple clinical measures of cholesterol associated with carriage of the e2 allele. As the unadjusted plots show (**Figure S3**), e2 homozygotes are dramatically different than other genotypes, though it should be noted that few e2 homozygotes were present in the population (n=16) and were within a limited age range (30-59 years). Selected lipid metabolites in the blood were positively associated with e2: a monoglyceride (MG) and four diacylglycerol (DG) metabolites (one of which was a replicate) were higher in e2 carriers compared to non-carriers. Diacylglycerol is a precursor to triacylglyceride (TG), which is typically higher in APOE2 carriers^24^. The effects of high DGs and TGs remains unclear. DG-rich diets fed to diabetic APOE-knockout mice had reduced atherosclerosis and lower plasma cholesterol than mice fed TG-rich or western diets^28,29^; however, non-targeted metabolomics studies have shown elevated levels of DGs and MGs in AD and MCI patient brains and blood compared to cognitively intact individuals^30,31^.

We observed six proteins that were significantly upregulated in APOE2 carriers (**Figure 3**). Though APOE2 is known to bind poorly to LDLR (~2% of e3 or e4 binding activity)^32^, APOE2 was associated with lower levels of LDL cholesterol across age groups as noted previously, perhaps due to compensatory up-regulation of LDL receptors (LDLR)^24^ (**Figure 3a**). APOE2 was associated with increased levels of the highly inducible HMOX-1, which has antioxidant properties and has been associated with neuroprotection and neurodegeneration^33^. SLAMF8 may be another link to an antioxidant effect of APOE2, as it has been implicated in modulation of reactive oxygen species and inflammation via negative regulation of NOX activity^34^. APOE2 carriers displayed higher levels of RNF31 (aka HOIP). HOIP is the catalytic component of the linear ubiquitin chain assembly complex (LUBAC), which was recently shown to have a role in the recognition and degradation of misfolded proteins^35^. Variation in CNTNAP2, a member of the neurexin superfamily of proteins involved in cell-cell interactions in the nervous system, has been associated with neurodevelopmental disorders^36^, and has been implicated in AD-related dementia^37^. Lastly, SRP14, which has a role in targeting secretory proteins to the rough endoplasmic reticulum (ER) membrane, has been identified as one of many tau-associated ER proteins in AD brains^38^.

**Figure 3.**
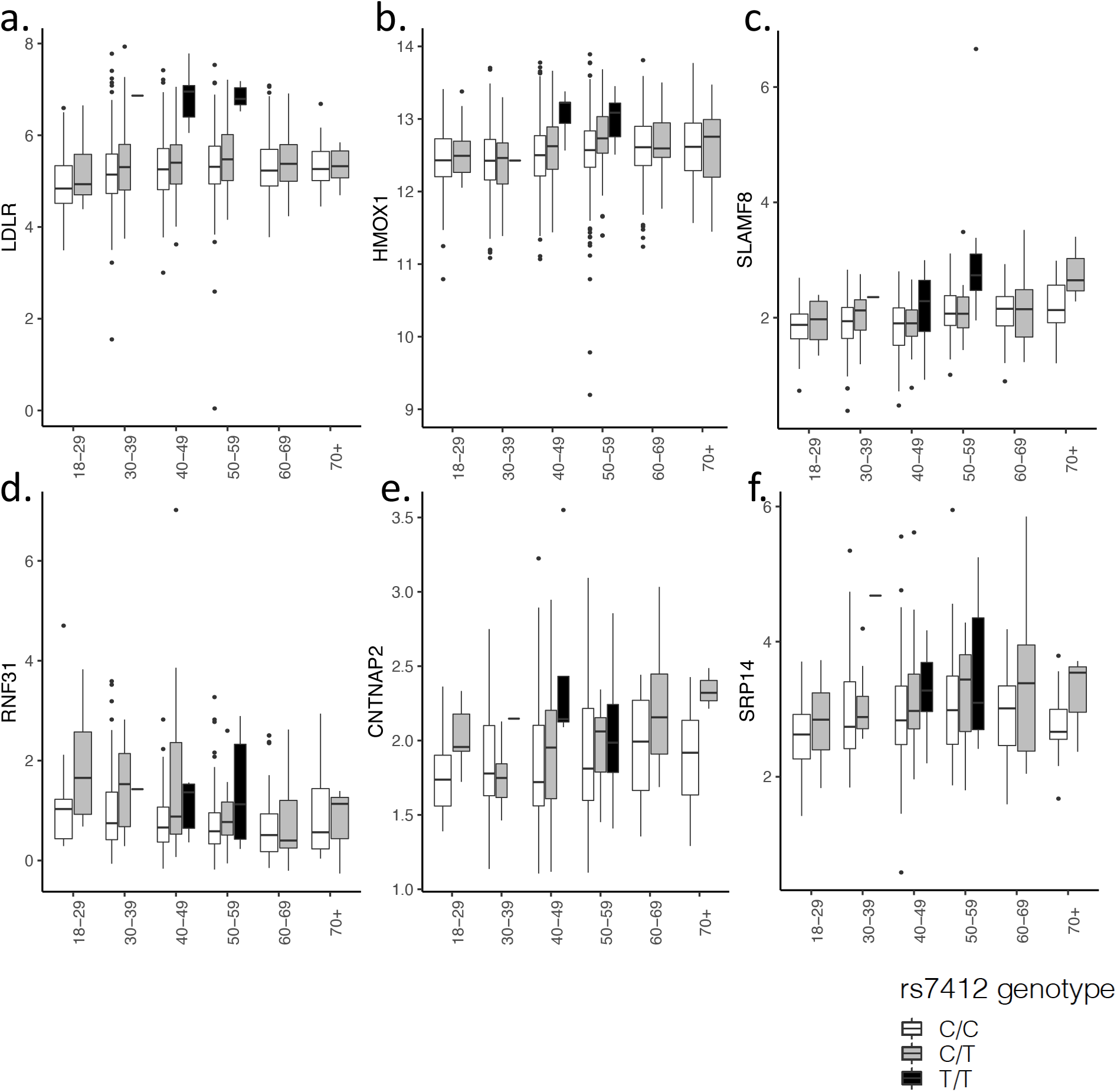
Unadjusted box plots of normalized protein expression levels (NPX) of six proteins significantly associated with APOE2 genotype, by age group. White boxplots=individuals who are homozygous for the major allele, gray boxplots=heterozygotes, black boxplots=minor allele homozygotes. Box plot midline=median value, lower/upper hinges=25^th^ and 75^th^ percentiles, respectively; lower whisker ends/upper whisker ends no further than 1.5 x interquartile range from the hinge. Data beyond whiskers are outlying points. **a.** Low-Density Lipoprotein Receptor (LDLR); **b.** heme oxygenase-1 (HMOX1); **c.** SLAM family member 8 (SLAMF8); **d.** E3 ubiquitin-protein ligase RNF31 (RNF31); **e.** Contactin-associated protein-like 2 (CNTNAP2); **f.** Signal recognition particle 14 kDa protein (SRP14).

#### ABCA7

*ABCA7* is involved in lipid efflux from cells into lipoprotein particles, plays a role in lipid homeostasis^39^, and has also been implicated in amyloid-β (Aβ) processing and deposition in the brain^40^. Our results support *ABCA7’s* lipid-related function by showing lower levels of two lactosylceramide (LC) metabolites among individuals carrying the AD-risk allele of rs3752246. These differences were evident across all ages but were especially pronounced in the youngest age groups (**Figure S4**). The minor allele of rs3752246 was also associated with higher levels of DEFA1 (neutrophil defensin 1, or alpha-defensin 1), an antimicrobial peptide. This novel gene-protein association is consistent with previous studies showing higher levels of this protein in CSF and sera of AD patients compared to controls^41,42^, potentially linking *ABCA7* with an inflammatory response pathway to AD.

#### INPP5D

An intronic SNP in *INPP5D* (rs10933431), which was associated with a lowered risk of AD in meta-analyses, was associated with lower levels of the protein IDUA (alpha-L-iduronidase) (**Figure S4**). *INPP5D,* which encodes the lipid phosphatase SHIP1, is a negative regulator of immune signaling and is expressed in microglia^43^. To our knowledge, this association has not been previously observed.

#### Polygenic risk score

No associations were observed between the *APOE*-free PGRS and any analyte after FDR correction for multiple testing, either in primary analyses or in analyses adjusted for e4 status, or among non-e4 individuals only. No effect modification by sex or APOE4 status was observed.

### Sex-specific findings

We observed a SNP x sex interaction involving the AD-protective *PICALM* variant, such that the minor allele was associated with higher levels of 30 proteins in men and lower levels of the proteins in women (**Figure 4, Figure S5**). These proteins were highly correlated with one another (mean pairwise spearman’s rho = 0.49); thus, it is unclear whether the associations are independently biologically meaningful, or whether there is a passenger effect, in which one or a few proteins are driving the sex-differential association with genotype observed in the data. The set of proteins that are differentially affected by sex and *PICALM* genotype are primarily implicated in immune processes, cell adhesion, and regulatory processes, with widely overlapping functions (**Figure S6**). In addition, the *PICALM* variant is associated with a sexspecific effect on five highly correlated long-chain fatty acid (LCFA) metabolites and one polyunsaturated fatty acid (PFA) metabolite (DHA) (**Figure 4**). A potential link between *PICALM* function, lipids, and AD is feasible: fatty acids, and DHA in particular, have long been known to have a role in maintaining brain health and cognition^44^, while *PICALM* expression has been shown to influence cholesterol homeostasis through multiple mechanisms^45^. To investigate further, we conducted a post-hoc analysis examining the impact of this variant on AD risk stratified by sex, in a meta-analysis of clinically diagnosed late-onset AD (18,812 individuals) (See Online Methods for details). While AD risk was reduced in both men and women among carriers of the minor allele, the effect was stronger among men (**Table 2 and Table S5**), which was consistent with the sex-stratified SNP-analyte analyses (data not shown).

**Figure 4.**
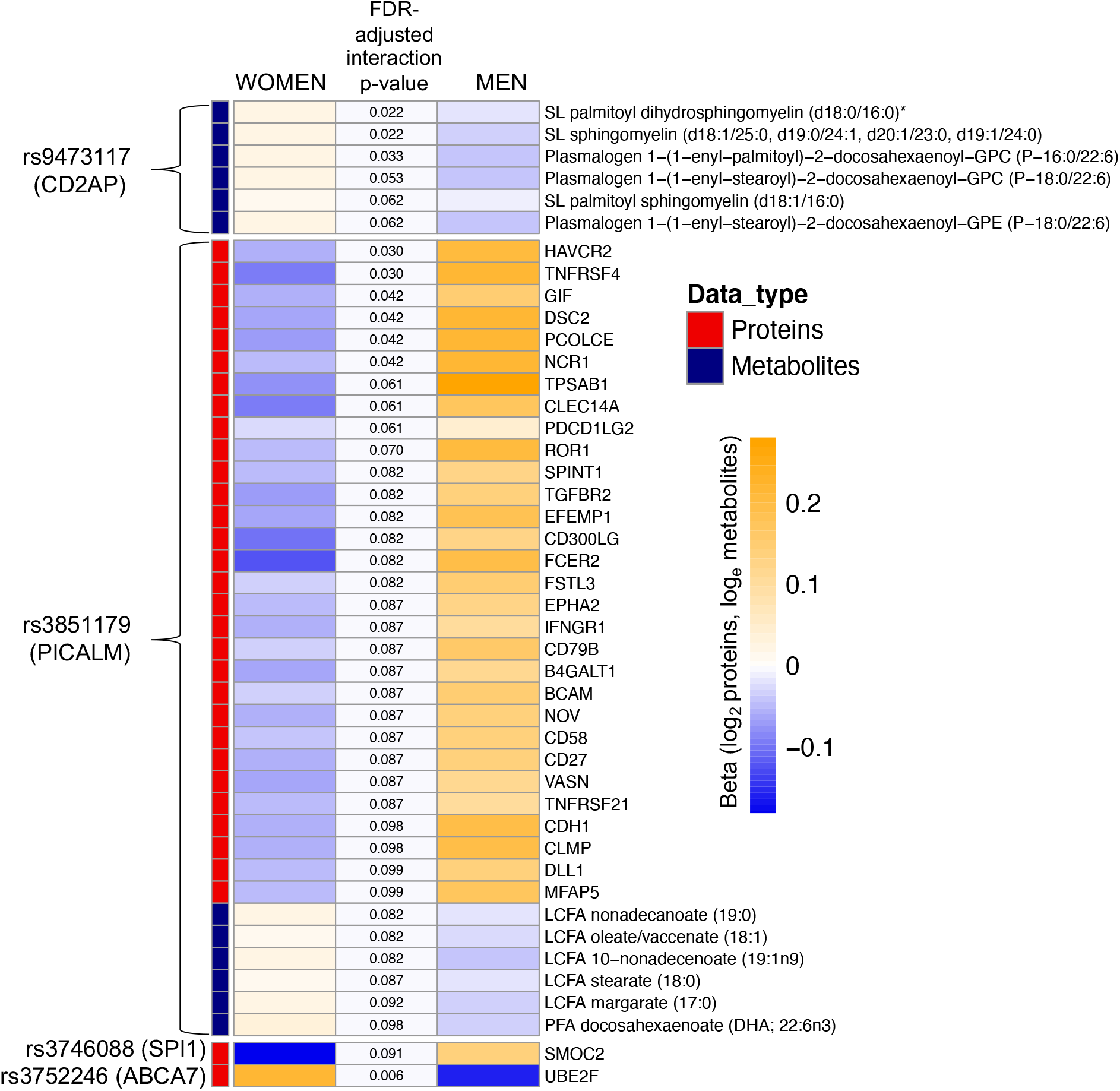
Heatmap of statistically significant genotype x sex interaction terms at FDR-adjusted p-value<0.1. Beta coefficients obtained from sex-stratified analyses, middle-column p-values from interaction term in the full model. SL=sphingolipid; LCFA=long-chain fatty acid; PFA=polyunsaturated fatty acid.

**Table 2.** Results of sex-specific analysis and sex-SNP interaction analysis of PICALM variant 3851179 in the ADGC^a^.

| **Sex-stratified results<sup>b</sup>** | **Beta** | **StdError** | **P-value** | **MAF** |
| --- | --- | --- | --- | --- |
| Male model 1 | -0.206 | 0.035 | 5.62E-09 | 0.358 |
| Male model 2 | -0.176 | 0.038 | 4.08E-06 | 0.359 |
| Female model 1 | -0.083 | 0.029 | 4.37E-03 | 0.354 |
| Female model 2 | -0.087 | 0.031 | 5.60E-03 | 0.352 |

| **Interaction Results<sup>c</sup>** | **Interaction beta** | **Std Error** | **P-value** | **MAF** |
| --- | --- | --- | --- | --- |
| Model 1 | 0.116 | 0.044 | 8.05E-03 | 0.354 |
| Model 2 | 0.372 | 0.048 | 7.84E-02 | 0.354 |

<sup>a</sup>N=9,135 cases (60% female), 9,677 controls (60% female)
<sup>b</sup>Model 1: adjusted for age, sex, and PCs; Model 2: adjusted for age, sex, PCs, and APOE genotype.
<sup>c</sup>Model 1: adjusted for age and PCs; Model 2: adjusted for age, PCs, and APOE.

Other observed sex-specific effects were more modest. The SNP near *CD2AP,* a scaffolding protein, interacted with sex to affect three highly correlated sphingomyelins and three plasmologens, while the SNP in *SPI1,* a transcription factor associated with microglial activation^46^, interacted with sex to affect SMOC2, a protein involved in microgliosis that has been previously associated with Aβ positivity in CSF^47^. Lastly, the missense *ABCA7* SNP interacted with sex to affect levels of Ubiquitin conjugating enzyme E2f (UBE2F).

### Stratification by self-identified race/ethnicity

Previous studies have shown genetic heterogeneity between white and non-white individuals, particularly with regard to African Americans and risk of cognitive outcomes among carriers of *APOE* and *ABCA7* variants^48,49^. Unfortunately, due to vanishingly small numbers in individual self-identified groups (**Table 1**), we were not able to assess genetic risk effects in individual groups with statistical rigor. As expected, analyses restricted to white individuals recapitulated results of the overall analysis (**Figure S7**). In the nonwhite group overall, we observed effect sizes that were consistent with the overall results and white-only results (**Figure S8**). Despite these overall consistencies, and given known wide-ranging racial/ethnic disparities in dementia incidence^50^, it is imperative that future deep-phenotyping studies are far more inclusive than the study presented here.

## Discussion

Our study examines associations between known genetic risk factors for AD and blood markers (clinical labs, proteins, and metabolites). It provides insight into the manifestation of AD-related genetic risk in blood-borne analytes from cognitively normal individuals and demonstrated how AD-related genetic variation manifests in the blood across adulthood. Our results contribute to the growing literature highlighting a potential causal variant (missense SNP in PILRA), point to new mechanisms of protection among APOE2 carriers, and suggest a role for infectious diseases as AD risk factors, alongside lipid metabolism, immune response, and endocytosis. We also uncovered intriguing differences between men and women in how genetic risk manifests in the blood. These analyses not only add to the existing literature on functional genomics in AD, but also offer up multiple potential new hypotheses to catalyze future studies.

The strongest associations in the study were between the *NYAP1* SNP (rs12539172) and the PILRB/PILRA proteins. PILRA and PILRB are paired inhibiting/activating receptors, respectively, that are expressed on innate immune cells, recognize certain O-glycosylated proteins, and have an important role in regulating acute inflammatory reactions^51^. The R78 substitution in PILRA (rs1859788) has been shown to reduce the binding capacity of endogenous ligands and thereby potentially increase microglial activity^20^. In addition, while controversial, work from our group and others^52–54^ has suggested a potential viral role in AD risk. Notably, the R78 variant has been implicated in HSV-1 infectivity^20^ and differences in HSV-1 antibody titer levels^17^. While previous studies have hypothesized that reduced activity of PILRA was due to steric conformational changes in the protein leading to reduced binding of key ligands (including HSV-1 glycoprotein B), our results suggest that reduced levels of circulating PILRA protein in R78 carriers could also be a factor in the overall protective effect of this genetic variant.

Strong associations were observed between multiple lipid analytes and the SNPs encoding APOE4 (rs429358) and APOE2 (rs7412). *APOE* normally plays a key role in lipid transport, including shuttling cholesterol to neurons in healthy brains. Notably, *APOE* has a role in Aβ metabolism, and while the exact mechanism is unknown, the e4 variant appears to accelerate neurotoxic Aβ accumulation, aggregation, and deposition in the brain^55^. Blood cholesterol levels among APOE4 and APOE2 carriers amongst Arivale participants were largely consistent with a large body of existing data, as this is well studied. It should be noted that while 5-10% of e2 homozygotes develop type III hyperlipoproteinemia (typically in the presence of an existing metabolic disorder^56^) resulting in elevated cholesterol levels, all e2 homozygotes in the study had significantly decreased levels of LDL cholesterol compared to other genotypes. Greater understanding of the compensatory mechanism leading to upregulated *LDLR* and lower circulating LDL cholesterol is needed. In addition, our results indicating a link between APOE2 and several potential protective mechanisms suggested by upregulation of HMOX1, SLAMF8, and RNF31 in e2 carriers, warrants further research.

Genetic variation likely affects men and women differentially, pointing to mechanisms that contribute to known differences in AD pathology between the sexes^57^. Our results highlight a novel interaction between the AD-risk variant in *PICALM* and multiple proteins implicated in immune response in a sex-specific manner, and support emerging research showing sex differences in the neuroimmune response that impact microglia function^58^. This multi-analyte interaction was supported by results from sex stratified GWAS meta-analyses, which showed differing effect sizes of the variant on men vs. women.

In addition to effects of individual genetic variants, we also examined an AD-specific polygenic risk score. While the PGRS is predictive of disease in case/control studies^59^, it was not associated with any blood analytes in the all-ages AD-free Arivale cohort. Combining genetic effects into a single score for AD likely served to dilute any individual genetic effect on the manifestation of genetic risk in the blood. In addition, the relative youth and cognitive health of this cohort should be taken into account. The PGRS may be more likely to detect perturbation in analytes that are markers of systemic inflammation or immune dysfunction in later life and among cohorts experiencing cognitive impairment.

The results presented here are novel and we believe will be of interest to the AD-related functional genomics community, though several limitations should be noted. The study population was not a random sample but was self-selected. The population is largely self-identified NHW, was mostly located on the west coast, and likely has higher than average socio-economic status (though these data were not captured). Thus, results may not be generalizable to a broader population. At this time, we were not aware of a suitable replication cohort that would contain parallel-omics panels in an all-ages health-heterogeneous cohort. Future studies will be needed to assess generality of the findings to other populations. Another limitation to the interpretation of results concerns the issue of pleiotropy; we cannot discern pleiotropic, non-AD-related effects from true causal effects that are implicated in AD pathogenesis. Related, we only obtained peripheral plasma, and are unable to examine effects in AD-relevant compartments such as brain or CSF. We only had high-coverage WGS available and did not interrogate other types of genetic variation such as copy number variants, indels, and short tandem repeats. Lastly, data harmonization with other studies will be a challenge. For instance, most previous metabolomics studies used metabolomics data that lacked complete speciation, and more work is needed with newer technologies that yield high fidelity data to determine the biological effects of specific serum metabolites.

This study also has multiple strengths. While most studies focused on AD-related genetic variation consist of case/control cohorts in older adults, the Arivale data offered an unprecedented look into how genetic variation perturbs physiological pathways in the blood long before disease onset, in health-heterogeneous individuals of all ages. This feature allowed us to observe subtle changes in blood associated with genetic variation, due to the relatively large sample size (2831 individuals with WGS) and the high quality of the blood analytes collected. Our results are from a “real-world” cohort, which promises to be an increasing source of large-scale data in the community going forward, with its accompanying advantages and disadvantages. Some results were previously unobserved and need to be replicated (such as the associations between *APOE2* and multiple proteins), while other results are in agreement with previous findings and serve to reinforce confidence that the results are reasonably representative and not simply spurious.

Due to a unified world-wide effort, dozens of genetic variants have been robustly implicated in the development of AD, though we are still in the early stages of understanding exactly how genetic variation contributes to disease. Our study showed that AD-related genetic variation manifests in the blood, from early adulthood onward, and highlights known targets for prevention in early and mid-life, such as cholesterol monitoring, mitigation of inflammation, and possibly, HSV-1 prevention and/or viral load management. Importantly, as well as yielding new insight into the pathobiology of AD through adulthood, these results may provide a significant number of new drug targets that are highly novel and biologically plausible or may serve as biomarkers if confirmed to have a consistent influence on AD pathophysiology. Lastly, these results highlight the need to assess for sex differences in future studies. Taken together, these results not only illustrate previously unobserved biological phenomenon implicated in development of AD, but also serve as an important resource for the generation of hypotheses for future functional genomics studies and emphasize the potential insight that can be gleaned from deeply phenotyped individuals.

## Online Methods

### Population

The Institute for Systems Biology, through partnership with their spin-out company Arivale, has access to a wealth of data collected longitudinally from subscribers in the commercially available Arivale Scientific Wellness program (as described previously^3,13^), starting in July 2015. In brief, participants in the Arivale program were assigned a health coach upon joining the program, who then utilized data from clinical blood assays and detailed health-history and behavioral questionnaires to personalize health advice and management of health goals; coaching generally focused on exercise, nutrition, stress management, and sleep. participants have consented to their de-identified data being used for research purposes.

### Blood-derived analytes

for each participant, fasting clinical blood laboratory tests were measured upon joining the program and at regular intervals for those enrolled longitudinally. Blood samples were collected at either local facilities hosted by LabCorp (North Carolina, USA) or Quest Diagnostics (New Jersey, USA). Whole genome sequencing was performed on DNA extracted from whole blood with library preparation using the Illumina TruSeq Nano Library prep kit and sequenced using Illumina HiSeq X, PE-150, target 30X coverage at a single CLIA-approved sequencing laboratory. At the baseline blood draw, 2827 of the 2831 individuals with sequenced whole genomes had up to 63 fasting clinical blood lab tests with no more than 20% missing. Clinical blood tests included standard markers for cardiometabolic health (lipid levels), diabetes, inflammation, kidney and liver function, nutrition (vitamins and minerals), and blood cell counts; blood samples were collected and processed by Quest Diagnostics and Labcorp. Frozen plasma samples were also sent to Olink (Olink Bioscience, Sweden) for targeted proteomics assays based on Olink’s proximity extension assay technique. Up to 2694 of these participants had quantitative proteomic data on 274 proteins from three Olink panels (Cardiovascular II (https://www.olink.com/products/cvd-ii-panel, Cardiovascular III (https://www.olink.com/products/cvd-iii-panel/), and Inflammation (https://www.olink.com/products/inflammation/). An additional 919 proteins (from 10 additional panels available at Olink at the time) were obtained from a subsample of 354 individuals, in which APOE e2/e2 and APOE e4/e4 genotypes were overrepresented. Since multiple batches were performed, previously generated pooled control samples were run with each batch and used for batch correction and multiple control samples were included on each plate. Olink provides protein measurements as “Normalized Protein Expression” (NPX) values, Olink’s arbitrary unit which is in Log2 scale. Up to 1855 of the participants had data from 754 metabolites. Aliquots of frozen plasma samples were sent to Metabolon, Inc. to conduct metabolomics assays using the Metabolon HD4 discovery platform. Relative concentration values were reported for each metabolite.

### SNP selection

We selected 25 common and somewhat-rare (>1% allele frequency) SNPs that were significantly associated with Alzheimer’s Disease in a recent large-scale meta-analysis based on updated data from the International Genomics of Alzheimer’s Project (IGAP)^5^. These variants have been widely studied, and are determined to be within or near genes in one of four pathways known to be associated with AD risk, including cholesterol metabolism, immune response, regulation of endocytosis, and protein ubiquination^5–8,60^. In addition to the variants replicated in Kunkle et al., 2019, we also included the SNP coding for APOE e2 (rs7412), as it is historically understudied due to its rarity and we hypothesized that associations with this SNP may yield valuable clues to the known protective effect of this genotype. The 25 SNPs were linked to 24 genes (two SNPs in APOE), as detailed in Table S1.

### Polygenic risk score calculation for AD

PGRS for age-associated AD risk was computed using summary statistics from the initial IGAP-driven GWAS meta-analysis^6^. Briefly, the set of SNPs included in the PGS was determined as follows. The Benjamini-Hochberg^15^ procedure was applied to the p-values for all SNPs tested in the GWAS to account for multiple testing by controlling the false discovery rate (FDR) at a 5% level. This FDR-filtered set of SNPs was then further pruned using linkage disequilibrium (LD): pairs of SNPs in close proximity capturing highly correlated information (r^2^ > 0.2) were identified, and the SNP with the smaller p-value in the pair was kept; this was repeated until all remaining SNPs were mutually uncorrelated (r^2^ < 0.2 for all pairs). The PGRS for each individual was then calculated by summing up the published effect size for each selected SNP multiplied by the number of effect alleles the individual carried for that SNP, across all of the selected SNPs. Missing genotypes were mean imputed using the effect allele frequency.

### Statistical analysis

The study population consists of all participants in the Arivale Wellness program with CLIA-laboratory-generated whole genome sequences. Following a phenome-wide association study approach (PheWAS) ^12,14^, the primary model for each SNP used linear regression, with genotype (0, 1, or 2, with 0 indicating homozygosity for the major allele and 2 indicating homozygosity for the minor allele) as the predictor, and each continuous quantitative analyte as the dependent variable. Clinical lab and metabolite values were natural log transformed to account for right skewness and outliers, with +1 added to each natural log transformation to prevent zero values. Proteomic quantities were presented as normalized protein expression, Olink’s arbitrary unit, which is in log2 scale. Genetic ancestry was represented by principal components (PCs) 1-4, calculated using previously described methods ^61^. All SNP models were adjusted for age, sex, genetic ancestry PCs 1-4, and vendor identification for the clinical labs. Secondary models tested effect modification by sex by including a gene x sex interaction term in the models. We accounted for multiple comparisons by applying the Benjamini-Hochberg method ^15^ at alpha=0.05 on a per-SNP basis and applied to the main effect of genotype in the primary models, while we set alpha=0.1 as the threshold for the gene x sex interaction models, as interaction terms are typically underpowered compared to main effects and we sought to fully explore sex effects for future hypothesis generation. The FDR rate took into account testing for all 2008 possible analytes, with the understanding that this adjustment was highly conservative given a high degree of correlation among multiple groups of analytes, and the fact that some analytes were sampled in only a subset of individuals. Both raw and adjusted p-values will be reported.

We also repeated the primary PheWAS approach with participants stratified by self-identified race, due to evidence for variable genetic risk for cognitive outcomes between white and nonwhite populations^48,49^. Differences may be due to gene-environment interactions impacting nonwhite populations as a result of sociocultural elements and/or structural inequities due to racism, and/or local ancestry-driven variation near specific loci. As the SNPs selected for this PheWAS were based on a meta-analysis consisting of non-Hispanic white (NHW) individuals, our results may be primarily driven by the fact that the majority of the Arivale population is also NHW, and thus results are not generalizable to other populations (we acknowledge that adjustment for genetic principal components may not be adequate to overcome such strong potential confounding factors). Unfortunately, due to small numbers of individuals in specific non-NHW racial and ethnic groups, which become vanishingly small when accounting for allele frequency and numbers of available samples (**Table 1**), we were not able to assess genetic risk effects in individual groups with statistical rigor and had to group all non-NHW participants into one stratum for analysis. The stratified NHW and nonwhite group analyses merely serve as an investigation into whether our primary results reflected the majority-NHW makeup of the Arivale population. PheWAS was applied as described above, with FDR to account for multiple comparisons.

In order to visualize genotype-analyte associations across adulthood, we created boxplots of the log-transformed analyte values by genotype, stratified by age group (by decade, from 18-29 to 70 and over). All statistical analyses were performed in R v3.5.1 (https://www.R-project.org/).

In post-hoc exploratory analysis focused on the SNP in the PICALM locus (rs3851179), sex-stratified and sex-interaction analyses was performed on 12,324 cases (57.7% female) and 11,453 controls (59.9% female) of European ancestry from the Alzheimer’s Disease Genetics Consortium (ADGC) (see Supplementary Table 4 for dataset details). Datasets were imputed to the Haplotype Reference Consortium (HRC)^62^ panel using the Michigan Imputation Server (https://imputationserver.sph.umich.edu/index.html#!). Standard pre-imputation quality control was performed on all datasets individually, including exclusion of individuals with low call rate, individuals with a high degree of relatedness, and variants with low call rate^63^. Individuals with non-European ancestry according to principal components analysis of ancestry-informative markers were excluded from the further analysis. Detailed descriptions of individual ADGC datasets can be found in Kunkle et al.^5^. Study-specific logistic regression analyses employed Plink^64^ for sex-interaction analysis and SNPTest^65^ for sex-stratified analysis. Sex-interaction, which analyzed the sex*variant interaction, and sex-stratified analysis of males and females separately, were performed for two separate models per analysis, one adjusting for age, sex and PCs (model 1) and a second adjusting for age, sex, PCs and *APOE* (model 2). Results were meta-analyzed with METAL using inverse variance-based analysis^66^.

## Author Contributions

L.Heath and N.D.P. conceived and designed the study; L.Heath, J.C.E., B.W.K., A.C.N, and E.R.M. performed data analyses and figure generation; A.T.M., J.C.L., B.W.K., E.R.M, N.E-T., L.Hood, N.R., and N.D.P. acquired the data; L.Heath, J.C.E., A.T.M., S.A.K., J.C.L., C.C.F., B.A.L., L.M.M., N.E-T., T.E.G., L.Hood, and N.D.P. made substantial contributions to interpretation of the results. L.Heath and N.D.P. were primary authors of the manuscript. All authors read and approved final manuscript.

## Data Availability

Qualified researchers can access the Arivale deidentified dataset supporting the findings in this study for research purposes. Requests should be sent to. The data are available to qualified researchers on submission and approval of a research plan.

## Code Availability

Code used for PheWAS statistical analysis is available through the Sage Bionetworks Github (https://github.com/Sage-Bionetworks/ADsnps_PheWAS.git).

## Acknowledgments

This study was supported by the National Institutes of Health, National Institute on Aging grants U01 AG046139 (N.E-T, T.E.G, N.D.P), R01 AG061796 (N.E-T), RF1 AG051504 (N.E-T), and R01-AG062634-01 (B.W.K, E.R.M).

ADGC. The National Institutes of Health, National Institute on Aging (NIH-NIA) supported this work through the following grants: ADGC, U01 AG032984, RC2 AG036528; Samples from the National Cell Repository for Alzheimer’s Disease (NCRAD), which receives government support under a cooperative agreement grant (U24 AG21886) awarded by the National Institute on Aging (NIA), were used in this study. We thank contributors who collected samples used in this study, as well as patients and their families, whose help and participation made this work possible; Data for this study were prepared, archived, and distributed by the National Institute on Aging Alzheimer’s Disease Data Storage Site (NIAGADS) at the University of Pennsylvania (U24-AG041689-01); NACC, U01 AG016976; NIA LOAD (Columbia University), U24 AG026395, U24 AG026390, R01AG041797; Banner Sun Health Research Institute P30 AG019610; Boston University, P30 AG013846, U01 AG10483, R01 CA129769, R01 MH080295, R01 AG017173, R01 AG025259, R01 AG048927, R01AG33193, R01 AG009029; Columbia University, P50 AG008702, R37 AG015473, R01 AG037212, R01 AG028786; Duke University, P30 AG028377, AG05128; Emory University, AG025688; Group Health Research Institute, UO1 AG006781, UO1 HG004610, UO1 HG006375, U01 HG008657; Indiana University, P30 AG10133, R01 AG009956, RC2 AG036650; Johns Hopkins University, P50 AG005146, R01 AG020688; Massachusetts General Hospital, P50 AG005134; Mayo Clinic, P50 AG016574, R01 AG032990, KL2 RR024151; Mount Sinai School of Medicine, P50 AG005138, P01 AG002219; New York University, P30 AG08051, UL1 RR029893, 5R01AG012101, 5R01AG022374, 5R01AG013616, 1RC2AG036502, 1R01AG035137; North Carolina A&T University, P20 MD000546, R01 AG28786-01A1; Northwestern University, P30 AG013854; Oregon Health & Science University, P30 AG008017, R01 AG026916; Rush University, P30 AG010161, R01 AG019085, R01 AG15819, R01 AG17917, R01 AG030146, R01 AG01101, RC2 AG036650, R01 AG22018; TGen, R01 NS059873; University of Alabama at Birmingham, P50 AG016582; University of Arizona, R01 AG031581; University of California, Davis, P30 AG010129; University of California, Irvine, P50 AG016573; University of California, Los Angeles, P50 AG016570; University of California, San Diego, P50 AG005131; University of California, San Francisco, P50 AG023501, P01 AG019724; University of Kentucky, P30 AG028383, AG05144; University of Michigan, P50 AG008671; University of Pennsylvania, P30 AG010124; University of Pittsburgh, P50 AG005133, AG030653, AG041718, AG07562, AG02365; University of Southern California, P50 AG005142; University of Texas Southwestern, P30 AG012300; University of Miami, R01 AG027944, AG010491, AG027944, AG021547, AG019757; University of Washington, P50 AG005136, R01 AG042437; University of Wisconsin, P50 AG033514; Vanderbilt University, R01 AG019085; and Washington University, P50 AG005681, P01 AG03991, P01 AG026276. The Kathleen Price Bryan Brain Bank at Duke University Medical Center is funded by NINDS grant # NS39764, NIMH MH60451 and by Glaxo Smith Kline. Support was also from the Alzheimer’s Association (LAF, IIRG-08-89720; MP-V, IIRG-05-14147), the US Department of Veterans Affairs Administration, Office of Research and Development, Biomedical Laboratory Research Program, and BrightFocus Foundation (MP-V, A2111048). P.S.G.-H. is supported by Wellcome Trust, Howard Hughes Medical Institute, and the Canadian Institute of Health Research. Genotyping of the TGEN2 cohort was supported by Kronos Science. The TGen series was also funded by NIA grant AG041232 to AJM and MJH, The Banner Alzheimer’s Foundation, The Johnnie B. Byrd Sr. Alzheimer’s Institute, the Medical Research Council, and the state of Arizona and also includes samples from the following sites: Newcastle Brain Tissue Resource (funding via the Medical Research Council, local NHS trusts and Newcastle University), MRC London Brain Bank for Neurodegenerative Diseases (funding via the Medical Research Council),South West Dementia Brain Bank (funding via numerous sources including the Higher Education Funding Council for England (HEFCE), Alzheimer’s Research Trust (ART), BRACE as well as North Bristol NHS Trust Research and Innovation Department and DeNDRoN), The Netherlands Brain Bank (funding via numerous sources including Stichting MS Research, Brain Net Europe, Hersenstichting Nederland Breinbrekend Werk, International Parkinson Fonds, Internationale Stiching Alzheimer Onderzoek), Institut de Neuropatologia, Servei Anatomia Patologica, Universitat de Barcelona. ADNI data collection and sharing was funded by the National Institutes of Health Grant U01 AG024904 and Department of Defense award number W81XWH-12-2-0012. ADNI is funded by the National Institute on Aging, the National Institute of Biomedical Imaging and Bioengineering, and through generous contributions from the following: AbbVie, Alzheimer’s Association; Alzheimer’s Drug Discovery Foundation; Araclon Biotech; BioClinica, Inc.; Biogen; Bristol-Myers Squibb Company; CereSpir, Inc.; Eisai Inc.; Elan Pharmaceuticals, Inc.; Eli Lilly and Company; EuroImmun; F. Hoffmann-La Roche Ltd and its affiliated company Genentech, Inc.; Fujirebio; GE Healthcare; IXICO Ltd.; Janssen Alzheimer Immunotherapy Research & Development, LLC.; Johnson & Johnson Pharmaceutical Research & Development LLC.; Lumosity; Lundbeck; Merck & Co., Inc.; Meso Scale Diagnostics, LLC.; NeuroRx Research; Neurotrack Technologies; Novartis Pharmaceuticals Corporation; Pfizer Inc.; Piramal Imaging; Servier; Takeda Pharmaceutical Company; and Transition Therapeutics. The Canadian Institutes of Health Research is providing funds to support ADNI clinical sites in Canada. Private sector contributions are facilitated by the Foundation for the National Institutes of Health (www.fnih.org). The grantee organization is the Northern California Institute for Research and Education, and the study is coordinated by the Alzheimer’s Disease Cooperative Study at the University of California, San Diego. ADNI data are disseminated by the Laboratory for Neuro Imaging at the University of Southern California. We thank Drs. D. Stephen Snyder and Marilyn Miller from NIA who are ex-officio ADGC members.

