## Supplementary material for "Manifestations of genetic risk for Alzheimer’s Disease in the blood: a cross-sectional multi-omic analysis in healthy adults aged 18-90+" (from another version of this preprint: version 1, 2021-03-28)

**Supplementary Table 1. Replicated candidate SNPs associated with AD in Kunkle et al., 2019, GWAS meta-analysis**

| SNP | Variant class | Nearest gene | Chromosome | MAF[a] | Major/ Minor Allele | OR[b] | 95% CI[b] | Meta-P[b] |
|---|---|---|---|---|---|---|---|---|
| rs4844610 | intronic | CRI1 | 1 | 0.16 | C/A | 1.17 | 1.13-1.21 | 3.6e-24 |
| rs6733839 | intronic | BIN1 | 2 | 0.39 | C/T | 1.20 | 1.17-1.23 | 2.1e-44 |
| rs10933431 | intronic | INPP5D | 2 | 0.27 | C/G | 0.91 | 0.88-0.94 | 3.4e-09 |
| rs9271058 | intergenic | HLA-DRB1 | 6 | 0.27 | T/A | 1.10 | 1.07-1.13 | 1.4e-11 |
| rs9473117 | intergenic | CD2AP | 6 | 0.26 | A/C | 1.09 | 1.06-1.12 | 1.2e-10 |
| rs12539172 | 3' UTR | NYAP1 | 7 | 0.30 | C/T | 0.92 | 0.90-0.95 | 9.3e-10 |
| rs10808026 | intronic | EPHA1 | 7 | 0.19 | C/A | 0.90 | 0.88-0.93 | 1.3e-10 |
| rs73223431 | intronic | PTK2B | 8 | 0.35 | C/T | 1.10 | 1.07-1.13 | 6.3e-14 |
| rs9331896 | intronic | CLU | 8 | 0.39 | T/C | 0.88 | 0.85-0.90 | 4.6e-24 |
| rs7920721 | upstream | ECHDC3 | 10 | 0.35 | A/G | 1.08 | 1.06-1.11 | 1.8e-11 |
| rs3740688 | intronic | SPI1 | 11 | 0.45 | T/G | 0.92 | 0.89-0.94 | 5.4e-13 |
| rs7933202 | intergenic | MS4A2 | 11 | 0.37 | A/C | 0.89 | 0.87-0.92 | 1.9e-19 |
| rs3851179 | intergenic | PICALM | 11 | 0.36 | C/T | 0.88 | 0.86-0.90 | 6.0e-25 |
| rs11218343 | intronic | SORL1 | 11 | 0.06 | T/C | 0.80 | 0.75-0.85 | 2.9e-12 |
| rs17125924 | intronic | FERMT2 | 14 | 0.09 | A/G | 1.14 | 1.09-1.18 | 1.4e-09 |
| rs12881735 | intronic | SLC24A4 | 14 | 0.21 | T/C | 0.92 | 0.89-0.94 | 7.4e-09 |
| rs593742 | intergenic | ADAM10 | 15 | 0.36 | A/G | 0.93 | 0.91-0.95 | 6.8e-09 |
| rs7185636 | intronic | IQCK | 16 | 0.20 | T/C | 0.92 | 0.89-0.95 | 2.4e-08 |
| rs62039712 | intergenic | WWOX | 16 | 0.10 | G/A | 1.16 | 1.10-1.23 | 3.7e-08 |
| rs138190086 | intergenic | ACE | 17 | 0.02 | G/A | 1.30 | 1.19-1.42 | 5.3e-09 |
| rs3752246 | missense | ABCA7 | 19 | 0.18 | C/G | 1.15 | 1.11-1.18 | 3.1e-16 |
| rs7412 | missense | APOE (ε2 allele) | 19 | 0.07 | C/T | *NA* | | |
| rs429358 | missense | APOE (ε4 allele) | 19 | 0.13 | T/C | 3.32 | (3.20-3.45) | 1.2e-881 |
| rs6024870 | intronic | CASS4 | 20 | 0.08 | G/A | 0.88 | 0.85-0.92 | 3.5e-08 |
| rs2830500 | intergenic | ADAMTS1 | 21 | 0.29 | C/A | 0.93 | 0.91-0.96 | 2.6e-08 |

[a]MAF in Arivale cohort

[b]Kunkle et al Meta-analysis results, except rs429358 (stage 1 results posted) and rs7412, which was not included in the previous study.

**Supplementary Table 2. AD-GWAS SNPs significantly associated with analytes in Arivale data (FDR p<0.05)**

| SNP | Analyte | Beta | SE | Un-adjusted p | FDR-adjusted p | Minor allele freq-uency | Sample N |
|---|---|---|---|---|---|---|---|
| ***SNPs associated with clinical chemistries*** | | | | | | | |
| rs429358 / APOE (e4) | LDL particle number | 0.094 | 0.013 | 2.84E-12 | 5.71E-09 | 0.13 | 2824 |
| rs429358 / APOE (e4) | LDL cholesterol | 0.071 | 0.012 | 2.03E-09 | 2.04E-06 | 0.13 | 2817 |
| rs429358 / APOE (e4) | Total cholesterol | 0.034 | 0.007 | 3.61-E06 | 1.81E-03 | 0.14 | 2825 |
| rs429358 / APOE (e4) | Triglyceride:HDL ratio | 0.071 | 0.016 | 9.67E-06 | 3.88E-03 | 0.14 | 2825 |
| rs429358 / APOE (e4) | HDL cholesterol | -0.044 | 0.010 | 2.25E-05 | 1.09E-02 | 0.14 | 2825 |
| rs429358 / APOE (e4) | Triglycerides | 0.065 | 0.017 | 1.41E-04 | 3.96E-02 | 0.14 | 2825 |
| rs7412 / APOE (e2) | LDL particle number | -0.221 | 0.018 | 5.19E-35 | 1.04E-31 | 0.07 | 2824 |
| rs7412 / APOE (e2) | LDL cholesterol | -0.160 | 0.016 | 2.95E-24 | 2.96E-21 | 0.07 | 2817 |
| rs7412 / APOE (e2) | Total cholesterol | -0.068 | 0.010 | 3.88E-12 | 2.60E-09 | 0.07 | 2825 |
| rs7412 / APOE (e2) | LDL small | -0.205 | 0.035 | 4.43E-08 | 1.78E-05 | 0.07 | 2824 |
| ***SNPs associated with proteins*** | | | | | | | |
| rs10933431 / INPP5D | IDUA (P35475) | -0.072 | 0.017 | 1.51E-05 | 3.02E-02 | 0.27 | 2691 |
| rs12539172 / NYAP1 | PILRB (Q9UKJ0) | -0.683 | 0.048 | 1.12E-36 | 2.24E-33 | 0.31 | 354 |
| rs12539172 / NYAP1 | PILRA (Q9UKJ1) | -0.326 | 0.033 | 2.29E-20 | 2.30E-17 | 0.31 | 354 |
| rs3752246 / ABCA7 | DEFA1 (P59665) | 0.149 | 0.034 | 1.24E-05 | 8.28E-03 | 0.17 | 353 |
| rs429358 / APOE (e4) | PLA2G7 (Q13093) | 0.159 | 0.030 | 1.94E-07 | 1.30E-04 | 0.23 | 353 |
| rs429358 / APOE (e4) | CD28 (P10747) | 0.091 | 0.024 | 1.41E-04 | 3.96E-02 | 0.23 | 354 |
| rs7412 / APOE (e2) | LDLR (P01130) | 0.240 | 0.035 | 7.39E-12 | 3.71E-09 | 0.07 | 2692 |
| rs7412 / APOE (e2) | HMOX1 (P09601) | 0.120 | 0.026 | 5.89E-06 | 1.69E-03 | 0.07 | 2691 |
| rs7412 / APOE (e2) | SLAMF8 (Q9P0V8) | 0.240 | 0.055 | 1.47E-05 | 2.80E-03 | 0.14 | 354 |
| rs7412 / APOE (e2) | RNF31 (Q96EP0) | 0.326 | 0.088 | 2.58E-04 | 4.28E-02 | 0.14 | 354 |
| rs7412 / APOE (e2) | CNTNAP2 (Q9UHC6) | 0.136 | 0.037 | 3.36E-04 | 4.43E-02 | 0.14 | 354 |
| rs7412 / APOE (e2) | SRP14 (P37108) | 0.287 | 0.079 | 3.53E-04 | 4.43E-02 | 0.14 | 354 |
| ***SNPs associated with metabolites*** | | | | | | | |
| rs3752246 / ABCA7 | Sphingolipid, N-nervonoyl-sphingosine (d18:1/24:1)* | -0.070 | 0.007 | 5.98E-21 | 1.20E-17 | 0.18 | 1884 |
| rs3752246 / ABCA7 | Sphingolipid, N-palmitoyl-sphingosine (d18:1/16:0) | -0.025 | 0.005 | 9.40E-07 | 9.43E-04 | 0.18 | 1895 |
| rs429358 / APOE (e4) | oleoyl-arachidonoyl-glycerol (18:1/20:4) [1]* | 0.043 | 0.012 | 1.90E-04 | 3.96E-02 | 0.14 | 1883 |
| rs429358 / APOE (e4) | palmitoyl-arachidonoyl-glycerol (16:1/20:4) | 0.069 | 0.019 | 2.01E-04 | 3.96E-02 | 0.14 | 1589 |
| rs429358 / APOE (e4) | oleoyl-arachidonoyl-glycerol (18:1/20:4) [2]* | 0.044 | 0.012 | 2.17E-04 | 3.96E-02 | 0.14 | 1873 |
| rs7412 / APOE (e2) | oleoyl-arachidonoyl-glycerol (18:1/20:4) [2]* | 0.075 | 0.016 | 4.80E-06 | 1.61E-03 | 0.07 | 1873 |
| rs7412 / APOE (e2) | oleoyl-arachidonoyl-glycerol (18:1/20:4) [1]* | 0.072 | 0.016 | 9.26E-06 | 2.32E-03 | 0.06 | 1883 |
| rs7412 / APOE (e2) | linoleoyl-arachidonoyl-glycerol (18:2/20:4) [1]* | 0.083 | 0.019 | 1.34E-05 | 2.80E-03 | 0.06 | 1872 |
| rs7412 / APOE (e2) | palmitoyl-arachidonoyl-glycerol (16:0/20:4) [2]* | 0.113 | 0.026 | 1.53E-05 | 2.80E-03 | 0.06 | 1593 |
| rs7412 / APOE (e2) | oleoyl-oleoyl-glycerol (18:1/18:1) [1]* | 0.057 | 0.016 | 2.78E-04 | 4.28E-02 | 0.06 | 1903 |
| rs7412 / APOE (e2) | Monoacylglycerol, 1-linoleoylglycerol (18:2) | 0.071 | 0.020 | 2.99E-04 | 4.28E-02 | 0.06 | 1888 |

#### Supplementary Table 3. SNP x sex interactions (interaction FDR p<0.1)

| SNP / Nearest Gene | Analyte | Interaction Beta | Un-adjusted $p_{int}$ | FDR-adjusted $p_{int}$ |
|---|---|---|---|---|
| ***SNP x Sex interactions associated with proteins*** | | | | |
| rs3851179 / PICALM | HAVCR2 (Q8TDQ0) | -0.257 | 1.64E-05 | 3.02E-02 |
| rs3851179 / PICALM | TNFRSF4 (P43489) | -0.307 | 3.00E-05 | 3.02E-02 |
| rs3851179 / PICALM | GIF (P27352) | -0.191 | 8.74E-05 | 4.22E-02 |
| rs3851179 / PICALM | DSC2 (Q02487) | -0.274 | 1.03E-04 | 4.22E-02 |
| rs3851179 / PICALM | PCOLCE (Q15113) | -0.282 | 1.12E-04 | 4.22E-02 |
| rs3851179 / PICALM | NCR1 (O76036) | -0.268 | 1.26E-04 | 4.22E-02 |
| rs3851179 / PICALM | TPSAB1 (Q15661) | -0.375 | 2.51E-04 | 6.08E-02 |
| rs3851179 / PICALM | CLEC14A (Q86T13) | -0.258 | 2.66E-04 | 6.08E-02 |
| rs3851179 / PICALM | PDCD1LG2 (Q9BQ51) | -0.077 | 2.73E-04 | 6.08E-02 |
| rs3851179 / PICALM | ROR1 (Q01973) | -0.251 | 3.50E-04 | 7.03E-02 |
| rs3851179 / PICALM | SPINT1 (O43278) | -0.168 | 4.54E-04 | 8.21E-02 |
| rs3851179 / PICALM | TGFBR2 (P37173) | -0.221 | 5.04E-04 | 8.21E-02 |
| rs3851179 / PICALM | EFEMP1 (Q12805) | -0.249 | 5.38E-04 | 8.21E-02 |
| rs3851179 / PICALM | CD300LG (Q6UXG3) | -0.234 | 7.39E-04 | 8.21E-02 |
| rs3851179 / PICALM | FCER2 (P06734) | -0.335 | 7.52E-04 | 8.21E-02 |
| rs3851179 / PICALM | FSTL3 (O95633) | -0.176 | 7.77E-04 | 8.21E-02 |
| rs3851179 / PICALM | EPHA2 (P29317) | -0.176 | 8.66E-04 | 8.65E-02 |
| rs3851179 / PICALM | IFNGR1 (P15260) | -0.170 | 9.05E-04 | 8.65E-02 |
| rs3851179 / PICALM | CD79B (P40259) | -0.192 | 1.01E-03 | 8.67E-02 |
| rs3851179 / PICALM | B4GALT1 (P15291) | -0.180 | 1.08E-03 | 8.67E-02 |
| rs3851179 / PICALM | BCAM (P50895) | -0.190 | 1.16E-03 | 8.67E-02 |
| rs3851179 / PICALM | NOV (P48745) | -0.199 | 1.17E-03 | 8.67E-02 |
| rs3851179 / PICALM | CD58 (P19256) | -0.169 | 1.20E-03 | 8.67E-02 |
| rs3851179 / PICALM | CD27 (P26842) | -0.203 | 1.28E-03 | 8.67E-02 |
| rs3851179 / PICALM | VASN (Q6EMK4) | -0.183 | 1.28E-03 | 8.67E-02 |
| rs3851179 / PICALM | TNFRSF21 (O75509) | -0.160 | 1.30E-03 | 8.67E-02 |
| rs3851179 / PICALM | CDH1 (P12830) | -0.248 | 1.61E-03 | 9.81E-02 |
| rs3851179 / PICALM | CLMP (Q9H6B4) | -0.271 | 1.66E-03 | 9.81E-02 |
| rs3851179 / PICALM | DLL1 (O00548) | -0.196 | 1.74E-03 | 9.94E-02 |
| rs3851179 / PICALM | MFAP5 (Q13361) | -0.216 | 1.78E-03 | 9.94E-02 |
| rs3740688 / SPI1 | SMOC2 (Q9H3U7) | -0.323 | 4.54E-05 | 9.12E-02 |
| rs3752246 / ABCA7 | UBE2F (Q969M7) | 0.396 | 2.77E-06 | 5.57E-03 |
| ***SNP x Sex interactions associated with metabolites*** | | | | |
| rs9473117 / CD2AP | SL palmitoyl dihydrosphingomyelin (d18:0/16:0)* | 0.039 | 8.96E-06 | 1.80E-02 |
| rs9473117 / CD2AP | SL sphingomyelin (d18:1/25:0, d19:0/24:1, d20:1/23:0, d19:1/24:0) | 0.060 | 5.76E-05 | 5.78E-02 |
| rs9473117 / CD2AP | SL palmitoyl sphingomyelin (d18:1/16:0) | 0.021 | 1.45E-04 | 9.72E-02 |
| rs9473117 / CD2AP | Plasmalogen 1-(1-enyl-palmitoyl)-2-docosahexaenoyl-GPE (P-16:0/22:6) | 0.054 | 1.97E-04 | 9.86E-02 |
| rs3851179 / PICALM | LCFA nonadecanoate (19:0) | 0.040 | 6.73E-04 | 8.21E-02 |
| rs3851179 / PICALM | LCFA oleate/vaccenate (18:1) | 0.043 | 6.91E-04 | 8.21E-02 |
| rs3851179 / PICALM | LCFA 10-nonadecenoate (19:1n9) | 0.059 | 7.01E-04 | 8.21E-02 |
| rs3851179 / PICALM | LCFA stearate (18:0) | 0.032 | 1.16E-03 | 8.67E-02 |
| rs3851179 / PICALM | LCFA margarate (17:0) | 0.050 | 1.42E-03 | 9.17E-02 |
| rs3851179 / PICALM | PUFA docosahexaenoate (DHA; 22:6n3) | 0.057 | 1.63E-03 | 9.81E-02 |

**Supplementary Table 4**. ADGC datasets meta-analyzed in sex-stratified and sex-interaction analysis of rs38511179.

| COHORT | Female_cases | Female_controls | Male_cases | Male_controls |
|---|---|---|---|---|
| ACT | 333 | 873 | 199 | 698 |
| ADC1 | 840 | 302 | 708 | 208 |
| ADC2 | 368 | 106 | 368 | 50 |
| ADC3 | 498 | 366 | 408 | 214 |
| ADC4 | 168 | 241 | 136 | 136 |
| ADC5 | 152 | 331 | 134 | 174 |
| ADC6 | 124 | 225 | 89 | 113 |
| ADC7 | 264 | 503 | 250 | 287 |
| ADNI | 113 | 70 | 155 | 103 |
| GSK | 379 | 455 | 287 | 257 |
| MAYO | 377 | 535 | 280 | 510 |
| MTV | 145 | 116 | 109 | 73 |
| OHSU | 82 | 84 | 50 | 69 |
| PFIZER | 374 | 412 | 322 | 350 |
| ROSMAP | 203 | 540 | 85 | 207 |
| TARC1 | 199 | 118 | 124 | 63 |
| TGEN2 | 433 | 177 | 235 | 188 |
| UKS | 342 | 87 | 254 | 83 |
| UMVUMSSM | 738 | 685 | 405 | 428 |
| UPITT | 790 | 525 | 465 | 304 |
| WASHU1 | 194 | 113 | 145 | 74 |
| | 7116 | 6864 | 5208 | 4589 |

**Supplementary Table 5. Full model results for rs3851179 by sex interaction analysis, and stratification by sex**

| Model | MarkerName | Allele1 | Allele2 | Freq1 | FreqSE | MinFreq | MaxFreq | Effect | StdErr | P-value | Direction | HetISq | HetChiSq | HetDf | HetPVal |
|---|---|---|---|---|---|---|---|---|---|---|---|---|---|---|---|
| **Interaction Results** | | | | | | | | | | | | | | | |
| Model 1 | 11:85868640_C_T | t | c | 0.3541 | 0.009 | 0.3352 | 0.3723 | 0.1157 | 0.0436 | 8.05E-03 | ++-++++++-+++++-+--++ | 0 | 16.404 | 20 | 0.6912 |
| Model 2 | 11:85868640_C_T | t | c | 0.3544 | 0.0091 | 0.3352 | 0.3723 | 0.0841 | 0.0478 | 7.84E-02 | ++-++++++-+++++-+---+ | 0 | 12.194 | 20 | 0.9092 |
| **Sex-Stratified Results** | | | | | | | | | | | | | | | |
| Male Model 1 | 11:85868640_C_T | t | c | 0.3581 | 0.0176 | 0.3193 | 0.3914 | -0.2058 | 0.0353 | 5.62E-09 | ---------------+----- | 0 | 17 | 20 | 0.6529 |
| Male Model 2 | 11:85868640_C_T | t | c | 0.3594 | 0.0171 | 0.3182 | 0.3852 | -0.1758 | 0.0382 | 4.08E-06 | ------------+--+-?--- | 0 | 14.811 | 19 | 0.7345 |
| Female Model 1 | 11:85868640_C_T | t | c | 0.3539 | 0.0148 | 0.3148 | 0.3976 | -0.0832 | 0.0292 | 4.37E-03 | ---+---++---+--+----- | 0 | 18.444 | 20 | 0.5582 |
| Female Model 2 | 11:85868640_C_T | t | c | 0.3525 | 0.0142 | 0.3005 | 0.3964 | -0.0868 | 0.0313 | 5.60E-03 | -----+--+---++-+----- | 0 | 15.938 | 20 | 0.7204 |

#### SUPPLEMENTARY FIGURE LEGENDS

**Supplementary Figure 1. Unadjusted box plots of clinical chemistries and metabolites significantly associated with APOE4 genotype, by age group.** White boxplots=individuals who are homozygous for the major allele, gray boxplots=heterozygotes, black boxplots=minor allele homozygotes. Box plot midline=median value, lower/upper hinges=25$^{th}$ and 75$^{th}$ percentiles, respectively; lower whisker ends/upper whisker ends no further than 1.5 x interquartile range from the hinge. Data beyond whiskers are outlying points. **a.** LDL particle number (log nmol/L); **b.** LDL cholesterol (log mg/dL); **c.** Total cholesterol (log mg/dL); **d.** Triglyceride:HDL ratio (log); **e.** Triglycerides (log mg/dL); **f.** HDL cholesterol (log mg/dL); **g.** NPX of T-cell-specific surface glycoprotein CD28 (CD28); **h.** Platelet-activating factor acetylhydrolase (PLA2G7); **i.** Steroid lipid cortisone.

**Supplementary Figure 2. LDL particle number among APOE4 genotypes, according to self-reported usage of cholesterol-lowering medications.** White boxplots=individuals who are homozygous for the major allele, gray boxplots=heterozygotes, black boxplots=minor allele homozygotes. Box plot midline=median value, lower/upper hinges=25$^{th}$ and 75$^{th}$ percentiles, respectively; lower whisker ends/upper whisker ends no further than 1.5 x interquartile range from the hinge. Data beyond whiskers are outlying points. Left panel shows individuals by genotype who had reported ever or currently using a cholesterol-lowering medication; right panel shows individuals who reported no use of a cholesterol-lowering medication (individuals who did not complete this assessment survey question were not included).

**Supplementary Figure 3. Unadjusted box plots of clinical chemistries and metabolites significantly associated with APOE2 genotype, by age group.** White boxplots=individuals who are homozygous for the major allele, gray boxplots=heterozygotes, black boxplots=minor allele homozygotes. Box plot midline=median value, lower/upper hinges=25$^{th}$ and 75$^{th}$ percentiles, respectively; lower whisker ends/upper whisker ends no further than 1.5 x interquartile range from the hinge. Data beyond whiskers are outlying points. **a.** LDL particle number (log nmol/L); **b.** LDL cholesterol (log mg/dL); **c.** Total cholesterol (log mg/dL); **d.** LDL small (log nmol/L); **e.** Diacylglycerol, oleoyl-arachidonoyl-glycerol (18:1/20:4); **f.** Diacylglycerol, linoleoyl-arachidonoyl-glycerol (18:2/20:4); **g.** dDacylglycerol, palmitoyl-arachidonoyl-glycerol (16:0/20:4); **h.** Monoacylglycerol, 1-linoleoylglycerol (18:2); **i.** Secondary bile acid, Glycodeoxycholate.

**Supplementary Figure 4. Unadjusted box plots of analytes significantly associated with ABCA7, HLA-DRB1, and INPP5D genotypes, by age group.** White boxplots=individuals who are homozygous for the major allele, gray boxplots=heterozygotes, black boxplots=minor allele homozygotes. Box plot midline=median value, lower/upper hinges=25$^{th}$ and 75$^{th}$ percentiles, respectively; lower whisker ends/upper whisker ends no further than 1.5 x interquartile range from the hinge. Data beyond whiskers are outlying points. **a.** Sphingolipid, N-nervonoyl-sphingosine (d18:1/24:1), by ABCA7 genotype; **b.** Sphingolipid, N-palmitoyl-sphingosine (d18:1/16:0), by ABCA7 genotype; **c.** Neutrophil defensin 1 (DEFA1), by ABCA7 genotype; **d.** C-C motif chemokine 19 (CCL19), by HLA-DRB1 genotype; **e.** Alpha-L-iduronidase (IDUA), by INPP5D genotype.

**Supplementary Figure 5. Unadjusted box plots of proteins with a significant interaction term between sex and PICALM genotype at FDR<0.1, by age group.** White boxplots=individuals who are homozygous for the major allele, gray boxplots=heterozygotes, black boxplots=minor allele homozygotes. Box plot midline=median value, lower/upper hinges=25$^{th}$ and 75$^{th}$ percentiles, respectively; lower whisker ends/upper whisker ends no further than 1.5 x interquartile range from the hinge. Data beyond whiskers are outlying points. For each protein, left panel shows protein levels by genotype in men; right panel shows protein levels by genotype in women.

**Supplementary Figure 6. Network indicating proteins with overlapping GO terms from PICALM-sex interaction analysis.** Figure completed in CLUEGO, a Cytoscape plug-in.

**Supplementary Figure 7**. **Statistically significant SNP-analyte associations in self-identified Non-Hispanic white individuals only, after correcting for multiple testing (threshold FDR-adjusted p-value=0.05), by SNP**. Top panel: log-transformed beta-coefficient from the linear regression model adjusted for sex, age, and genetic principal components 1-4; markers above the zero line (orange) indicate analytes that increased in value with the minor allele, while markers below the line indicate markers that decreased in value. Second panel: FDR-adjusted $-\log_{10}$ p-value; orange line at FDR-p=0.05. Proteins=red, metabolites=blue, clinical chemistries=purple. Metabolite codes: DG=diacylglycerol; LC=lactosylceramide; o=oleoyl; a=arachidonoyl; g=glycerol; l=linoleoyl; p=palmitoyl. Third panel: minor allele frequency (MAF). Bottom panel: Total sample size for each analyte-SNP regression.

**Supplementary Figure 8**. **Statistically significant SNP-analyte associations in individuals who did not self-identify as non-Hispanic white only, after correcting for multiple testing (threshold FDR-adjusted p-value=0.05), by SNP**. Top panel: log-transformed beta-coefficient from the linear regression model adjusted for sex, age, and genetic principal components 1-4; markers above the zero line (orange) indicate analytes that increased in value with the minor allele, while markers below the line indicate markers that decreased in value. Second panel: FDR-adjusted $-\log_{10}$ p-value; orange line at FDR-p=0.05. Proteins=red, metabolites=blue, clinical chemistries=purple. Metabolite codes: DG=diacylglycerol; LC=lactosylceramide; o=oleoyl; a=arachidonoyl; g=glycerol; l=linoleoyl; p=palmitoyl. Third panel: minor allele frequency (MAF). Bottom panel: Total sample size for each analyte-SNP regression.

Supplementary Figure 1.

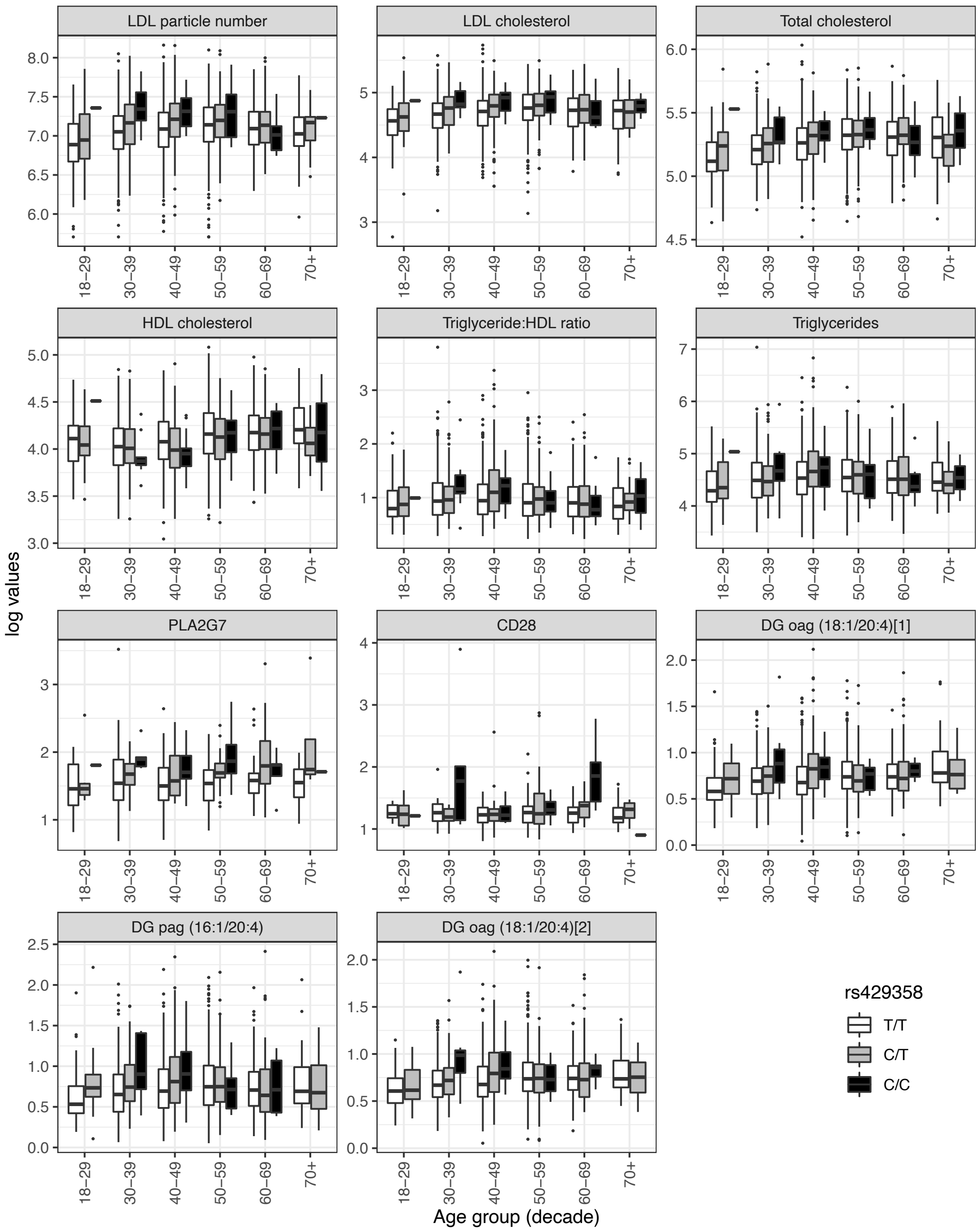

Supplementary Figure 2. LDL particle number among APOE4 genotypes, according to self-reported usage of cholesterol-lowering medications

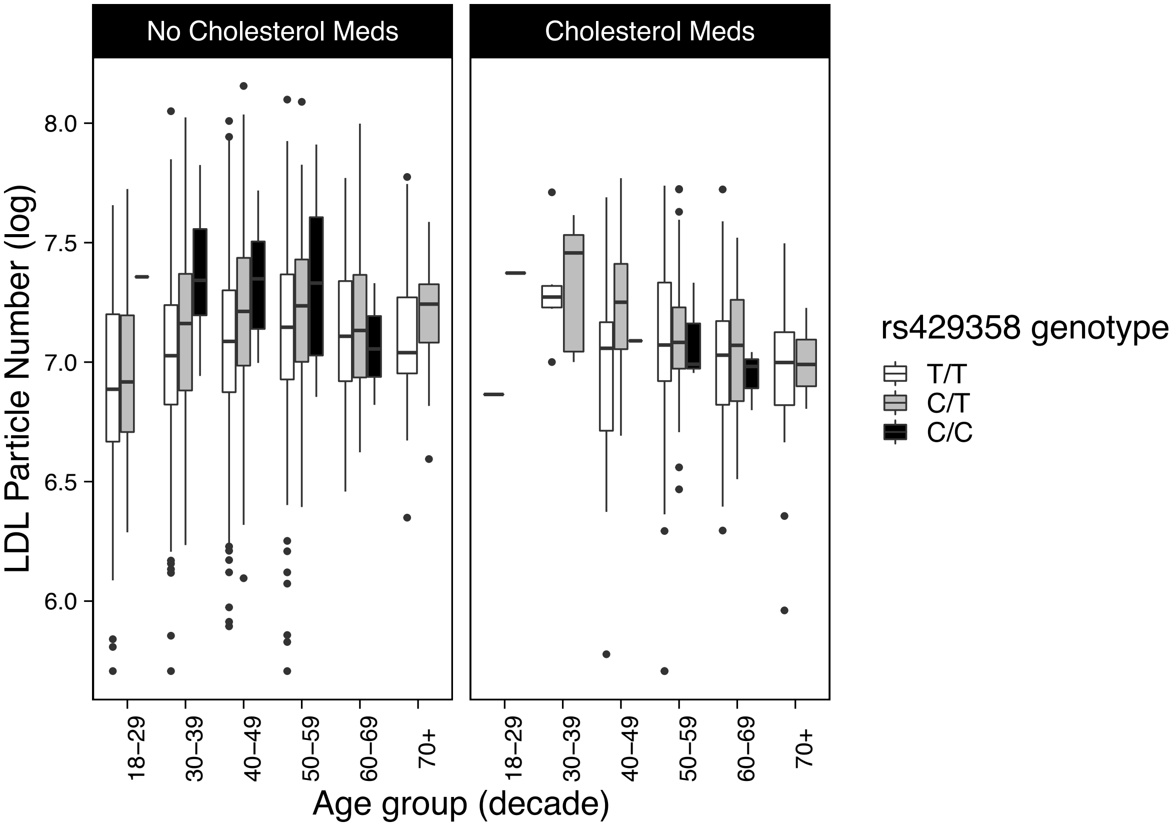

Supplementary Figure 3.

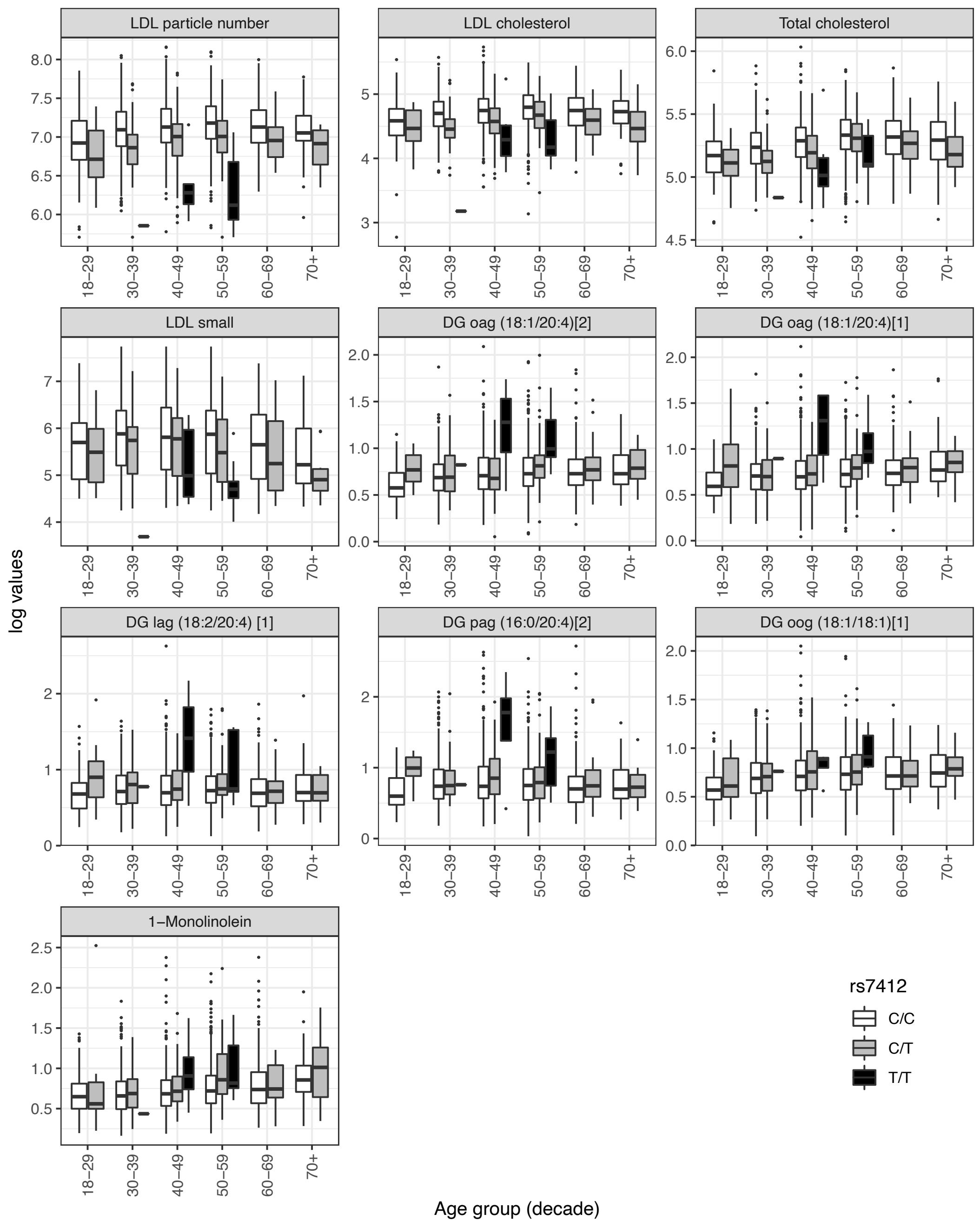

### Supplementary Figure 4.

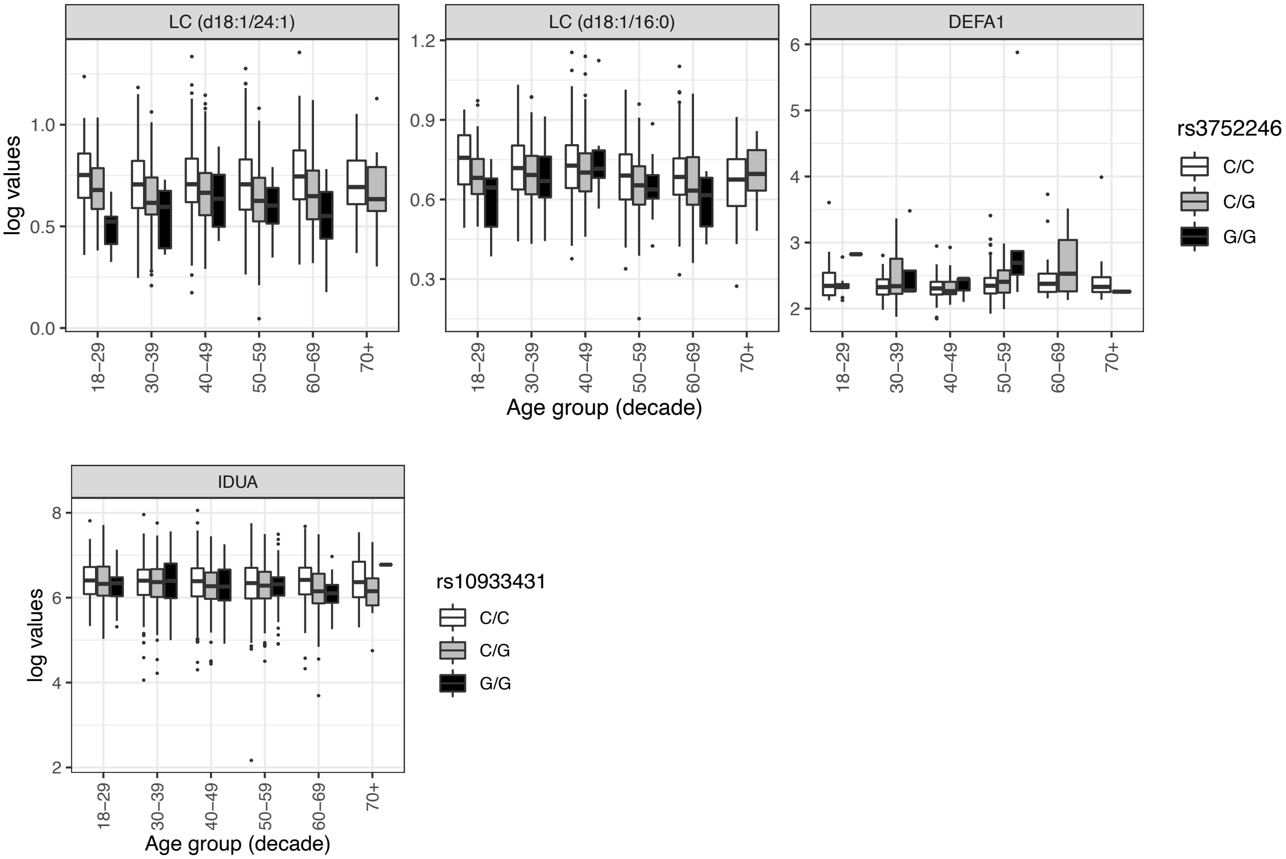

Supplementary Figure 5.

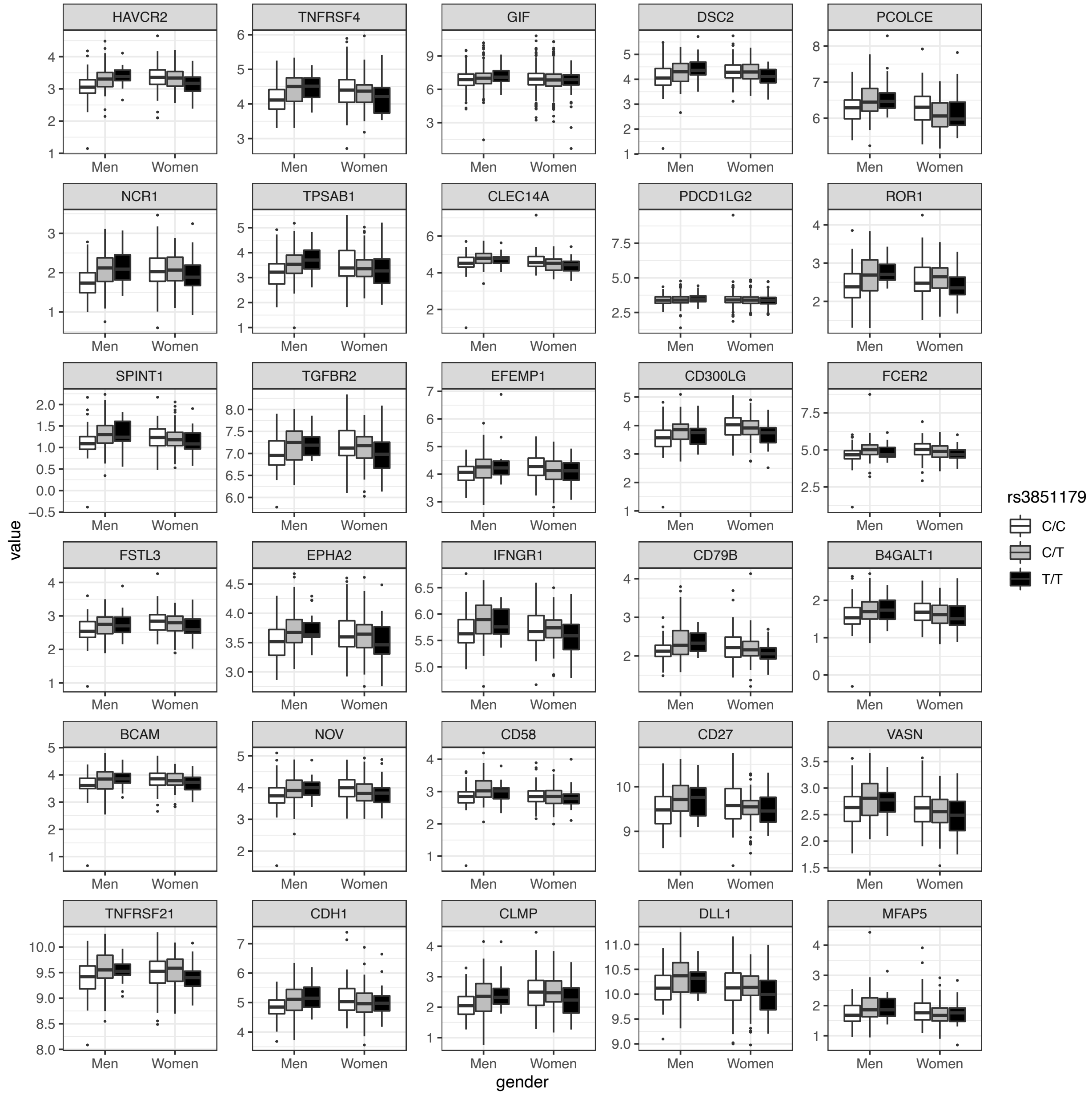

Supplementary Figure 6.

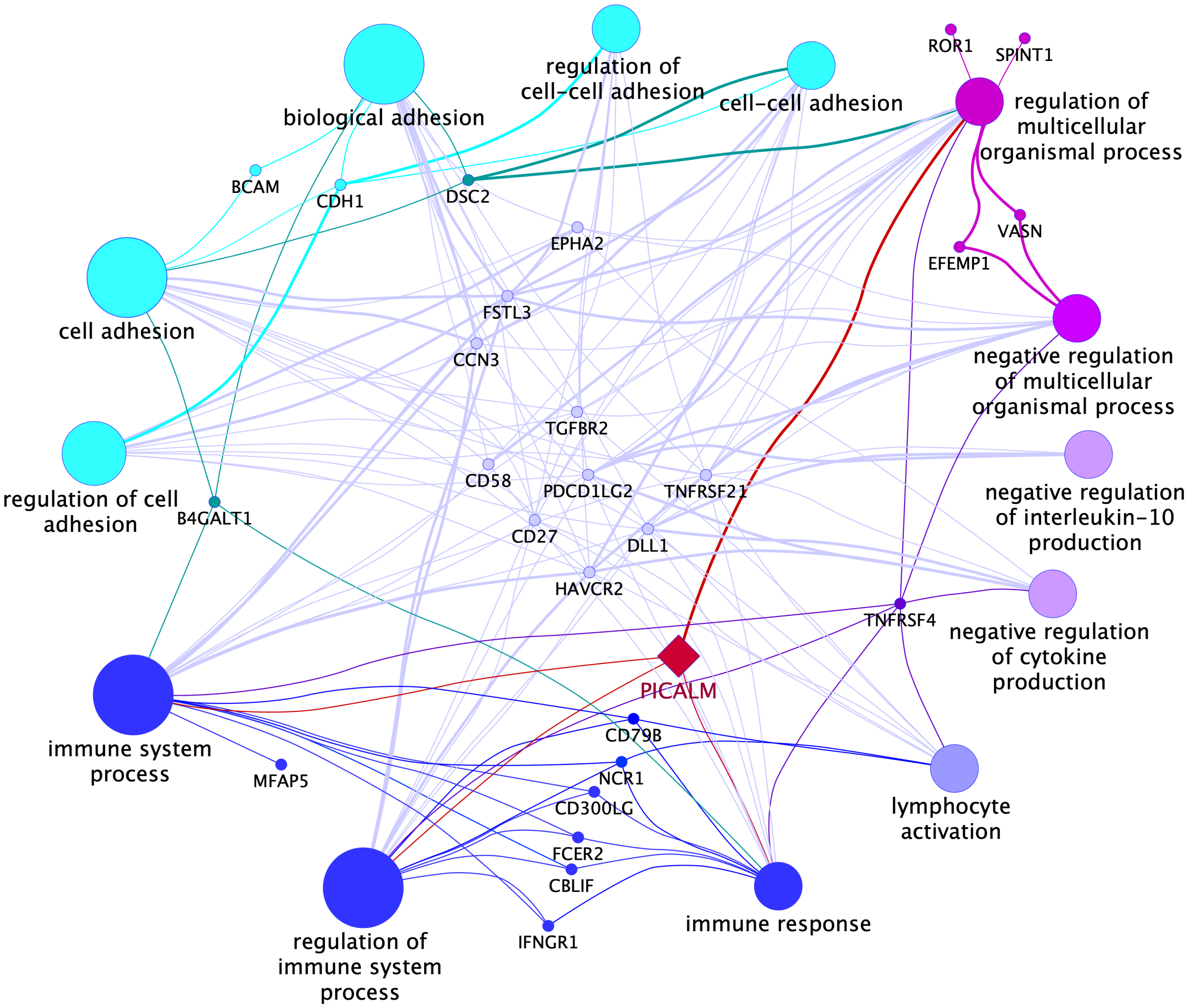

Supplementary Table 7. Non-Hispanic whites only

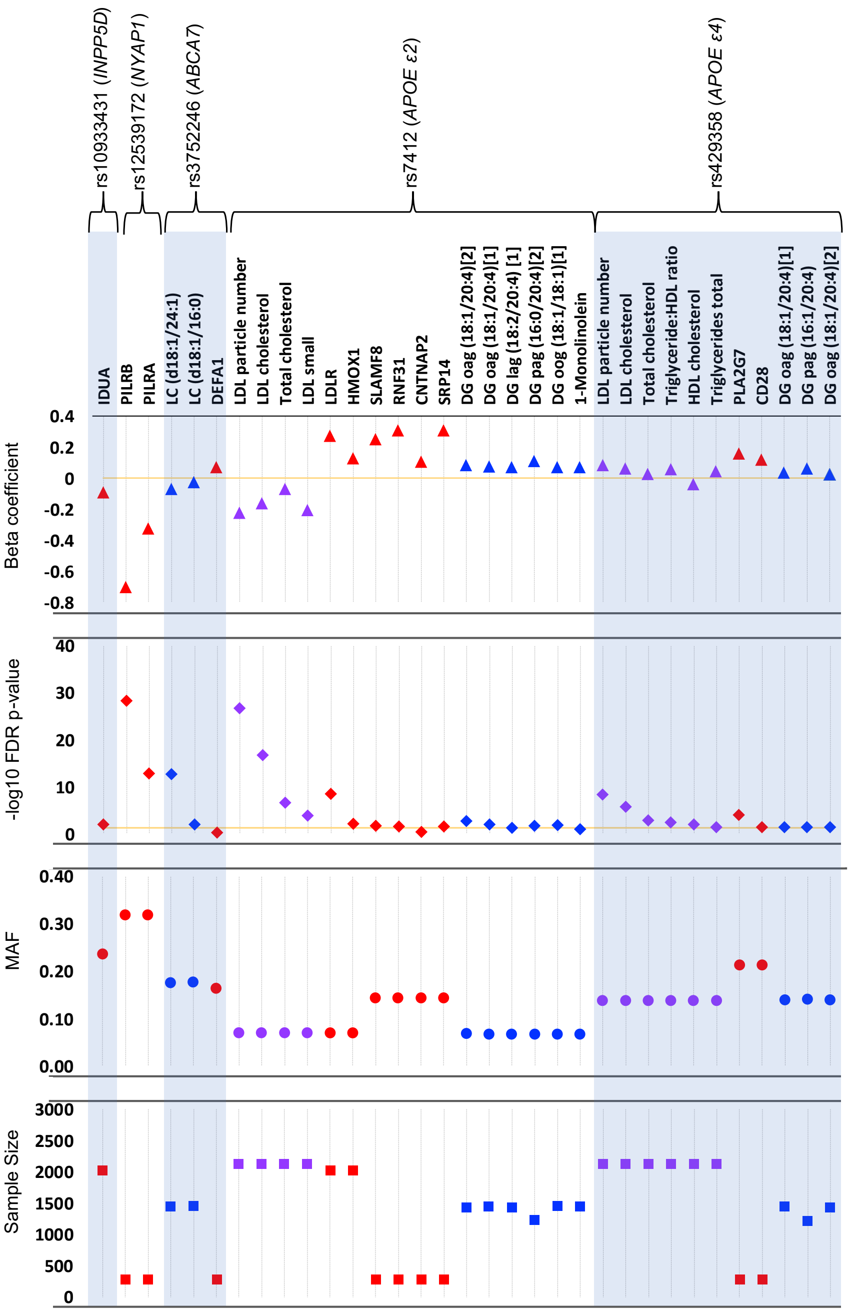

### Supplementary Table 8. Non-white only

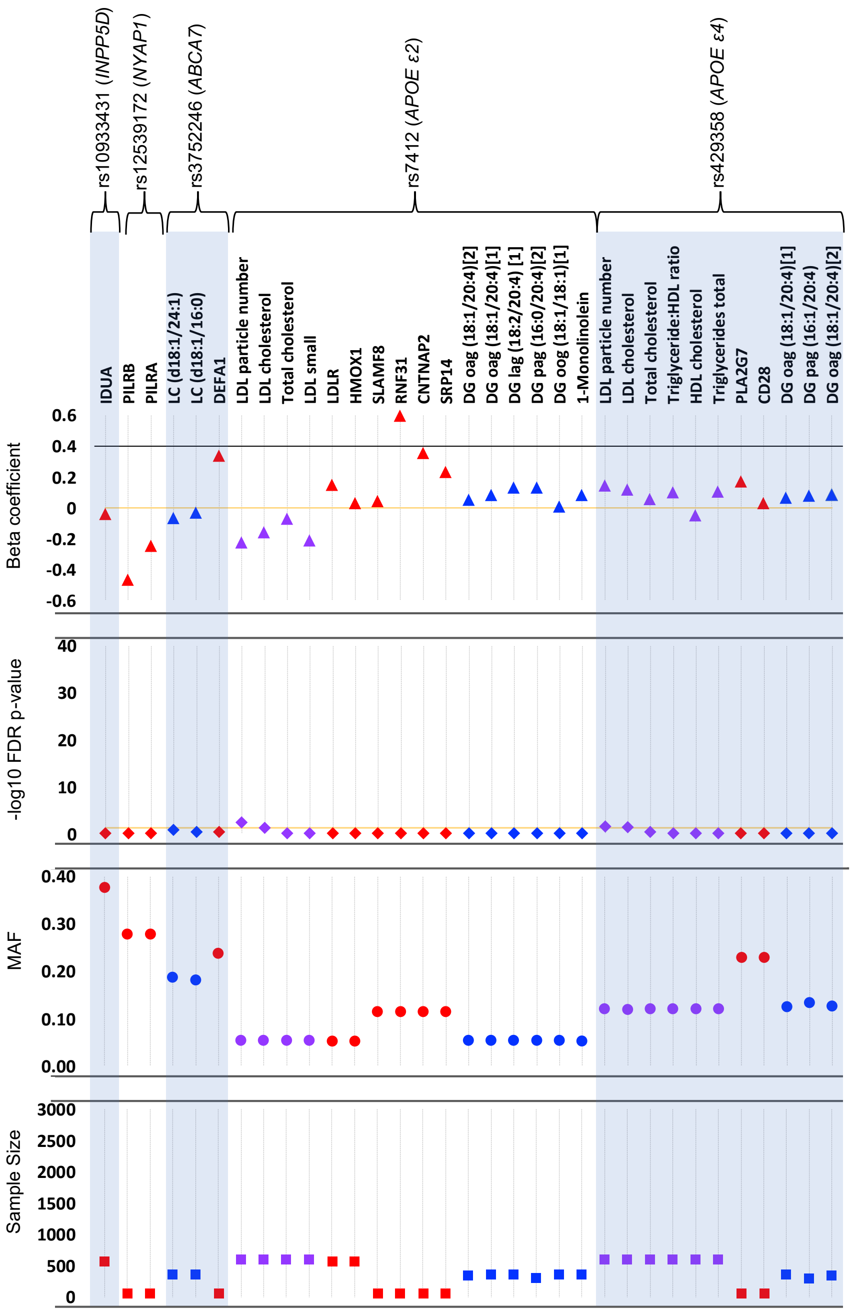
